# Assembly and architecture of the type III secretion sorting platform

**DOI:** 10.1101/2022.09.23.509200

**Authors:** J. Eduardo Soto, Jorge E. Galán, María Lara-Tejero

## Abstract

Type III secretion systems are bacterial nanomachines specialized in protein delivery into target eukaryotic cells. The structural and functional complexity of these machines demand highly coordinated mechanisms for their assembly and operation. The sorting platform is a critical component of type III secretion machines that ensures the timely engagement and secretion of proteins destined to travel this export pathway. However, the mechanisms that lead to the assembly of this multi-component structure have not been elucidated. Herein, employing structure modeling and an extensive *in vivo* crosslinking strategy, we provide a detailed inter-subunit-contact survey of the entire sorting platform complex. Using the identified crosslinks as signatures for pairwise inter-subunit interactions in combination with systematic genetic deletions, we mapped the assembly process of this unique bacterial structure. Insights generated by this study could serve as the bases for the development of anti-virulence strategies to combat several medically important bacterial pathogens.

## INTRODUCTION

Protein secretion mechanisms are central for the virulence of most bacterial pathogens. Through the activities of proteins secreted by a variety of protein secretion machines, bacterial pathogens can colonize the host, modulate host-cellular functions, and induce pathology^1^. One such protein secretion machines is the type III protein secretion system (T3SS), which has specifically evolved not just to secrete effector proteins from bacterial cells but to deliver them directly into target eukaryotic cells^2-5^. These effector proteins hijack host molecular functions to ensure bacterial survival and replication^6-8^. The adaptive advantage conferred by T3SSs is reflected by their broad phylogenetic distribution across diderm-LPS bacteria, and the variety of ecological niches where these machines are employed, ranging from animal and plant pathogens to fungi and protozoa symbionts^9^. *Salmonella enterica* serovar Typhimurium (*S*. Typhimurium), a worldwide cause of gastroenteritis disease, employs two T3SSs encoded within its so-called *Salmonella* pathogenicity islands 1 (SPI-1) and 2 (SPI-2), which are critical for its virulence^7,10^.

From a structural point of view, the core component of the entire T3SS structure or injectisome is the needle-complex, composed of a multi-ring base spanning both bacterial membranes, which serves as a conduit to enable effector proteins to get across the bacterial envelope^11,12^. Anchored to this base, is a hollow extracellular needle-like structure that provides a continuous channel for direct passage of effector proteins into the target host cell. The needle complex is in turn functionally powered by the export apparatus^13,14^, a set of inner membrane proteins enclosed within the base, and by a large cytosolic complex known as the sorting platform^15^, that plugs into the cytoplasmic portion of the needle complex. The T3SSs share a common evolutionary history with the bacterial flagella. Therefore, many of its core structural features are conserved between both biological machines including the entire export apparatus and some of the inner membrane and cytosolic components^16^.

Due to their high degree of structural complexity, assembly of T3SS machines requires exquisite coordination between multiple membrane, cytoplasmic, and extracellular building blocks. The biogenesis of the secretion machine is initiated at the inner bacterial membrane with the deployment of the export apparatus, a conical pseudo-helical structure that templates the assembly of the multiring membrane-spanning base^13,17^. The base is composed of a 15-mer of the InvG (SctC in the unified nomenclature, see Supplementary Table S1) secretin at the outer membrane (OM), and two nested rings at the inner membrane (IM). The inner and outer rings are formed by 24 subunits of the lipoprotein PrgK (SctJ), and the single-pass membrane protein PrgH (SctD), respectively. These stacked IM rings enclose the export apparatus at its center. Finally, by an unknown mechanism, the cytoplasmic sorting platform is recruited by the cytoplasmic N-terminal domain of PrgH^18^. The assembly of the sorting platform marks the onset of type III protein secretion. Escorted by dedicated chaperones and aided by regulatory proteins, substrates are sequentially loaded onto the sorting platform in a hierarchical fashion: first early substrates comprising the subunits that form the inner rod and the extracellular needle, followed by the translocases that facilitates passage of the effector proteins through the target host-cell membrane, and lastly the effector proteins that modulate cellular functions^15,19^.

The needle complex from different bacterial species has been amenable to purification and single particle cryo-electron microscopy (cryo-EM) analysis, which has resulted in high-resolution structures of all its components^11,20,21^. In contrast, the sorting platform has remained refractory to isolation as it is detached during the purification process. Recent advances in cryo-electron tomography (cryo-ET) have bypassed the need to isolate protein complexes, allowing the direct visualization of the intact T3SS machine in its native cellular environment. Cryo-ET imaging of the *Salmonella*^18^ and *Shigella*^22^ injectisomes revealed an overall conserved architecture for the sorting platform, resembling a hexagonal cage composed of six equidistant pods radially arranged about a central core. The cage-like structure is capped at its cytoplasmic end by a six-spoke structure or “cradle”. By adding traceable densities to individual components of the *S*. Typhimurium sorting platform, it has been possible to grossly assign the density maps to single proteins^18^. These studies have shown that each of these six pods are formed by OrgA (SctK) at their most membrane-proximal portion, and by SpaO (SctQ) at their distal section, which is responsible for the bulk of the protein densities associated with these structures. The spoke-like structure is constituted by OrgB (SctL), which emerges from the end of each of the SpaO pods and forms the cradle that serves as a platform for the hexameric InvC (SctN) ATPase. The ATPase is crowned at its top by the central stalk protein InvI (SctO), which in turn couples the ATPase function to the export gate major protein InvA (SctV). A unique structural feature revealed by the cryo-ET imaging is that the apparent symmetry disparity between the assembled inner membrane ring (24-fold) and the sorting platform (6-fold) is alleviated by the reorganization of the otherwise continuous PrgH_N_ cytoplasmic ring into six discrete tetrameric patches, onto which the six-pod cage-like sorting platform ultimately docks^18^, although the molecular mechanisms behind this striking rearrangement are unknown.

Despite the advances in the understanding of the overall shape and the gross topology of the sorting platform, the structural and mechanistic bases that govern its assembly are still poorly understood. Moreover, the lack of high-resolution structures of the cytosolic sorting platform that could define the interacting surfaces of its components has hampered structure-function studies. Here we generated a comprehensive and detailed topological map of the entire sorting platform using structural modeling and quantitative *in vivo* photo-crosslinking. Employing the identified crosslinked complexes as reporters for inter-subunit interactions, in combination with genetic analysis, we were able to delineate the assembly pathway that controls the formation of the *Salmonella* sorting platform.

## Results

### Defining the OrgA-PrgH interaction network

*In situ* cryo-ET and genetic analyses studies have postulated that the SPI-1 T3SS protein OrgA is the *Salmonella* sorting platform component that links the pod structures to the envelope-embedded needle complex^18^. However, direct experimental evidence for this notion is lacking. Furthermore, the absence of atomic information about the putative arrangement of OrgA and its direct interacting partners, has prevented the definition of the relevant interacting surfaces, which is crucial to enable the detail study of the sorting platform assembly process. To help clarify this issue we carried out a deep-mutagenesis screen to identify OrgA loss-of-function mutants. We reasoned that, mutations in amino acids involved in the formation of relevant functional interfaces between OrgA and other components of the sorting platform, should be enriched among mutations resulting in a loss-of-function phenotype. To search for these mutants at large-scale and in an unbiased fashion, we measured the functional effects of hundreds of mutations in parallel by deep mutational scanning^23^. We used error-prone PCR to generate a plasmid library of point mutants in *orgA*, which were then screened for their ability to invade cultured epithelial cells, a phenotype strictly dependent on the function of the SPI-1 T3SS. Next-generation sequencing was used to compare the frequency of each *orgA* mutant variant in the initial (input) and in the selected (output) libraries, and the log_2_-scaled enrichment ratio was calculated to generate a fitness score of each variant (Fig S1A). This screen allowed us to identify several point mutants with a significant negative effect on OrgA function (Fig S1B and S1C). However, further analysis of the resulting mutants indicated that, rather than clustering around specific domains of OrgA, the mutants were distributed all along the length of its sequence (Fig. S1C), suggesting that most of the mutations may have resulted in gross conformational changes in the OrgA structure. Furthermore, the lack of enrichment in specific domains of OrgA may also suggest that its interacting surfaces are most likely redundant so that changes in single amino acids may not result in strong phenotypes. We therefore concluded that this approach would not be well suited for the specific identification of OrgA amino acids directly involved in inter-subunit interactions.

To address the observed limitations of the genetic analysis and to identify OrgA amino acids directly involved in its interactions with other T3SS structural components we used a site-specific photo-crosslinking strategy. This approach involves the use of the photo-crosslinkable amino acid *p*-benzoyl-L-phenylalanine (*p*Bpa) that is incorporated into the protein of interest with the assistance of an orthogonal aminoacyl-tRNA synthetase/tRNA pair, which in turn suppresses the amber codon TAG^24^. Since the defined reactive radius of *p*Bpa is ∼3.1 Å^25^, this experimental approach allowed us to map the interfaces between OrgA and other components of the sorting platform in an *in vivo* context and at residue-level resolution. To guide the proper placement of *p*Bpa within the interacting protein surfaces we obtained structural insights into the conformation of the sorting platform pods by leveraging the predictive power of the recently released AlphaFold 2 (AF2)^26^, a deep learning-based algorithm that performs predictions of structures of proteins or protein complexes^27^. Based on our previous cryo ET studies, which have shown that the cytoplasmic domain of PrgH (PrgH^1-130^) and OrgA form the proximal region of the sorting platform pods, we modeled a complex made up of two molecules of PrgH^1-130^ and one molecule of OrgA. We then used the predicted structural information to introduce *p*Bpa within the predicted interacting surfaces. The model predicts that OrgA interacts with the cytoplasmic domain of two molecules of PrgH, each oriented differently from one another (Fig. 1A). As a result, in this model each of the two PrgH molecules engages OrgA in a different fashion although both do so through their forkhead-associated (FHA) domain (Fig. S2). This domain typically consists of a β-sandwich domain formed by two β-sheets interconnected by inter-strand loops. The loops located at the apical face of the FHA domains typically confer specificity for phospho-threonine-containing proteins^28^. However, in the case of PrgH, the residues responsible for phosphopeptide recognition are not conserved^29^, suggesting that the FHA domain of this T3SS component recognizes its binding partners in a phospho-independent manner. Based on the predicted model, we introduced the unnatural amino acid *p*Bpa at 10 positions between the β-strands of the FHA domain of PrgH (Fig. 1A and S2). All pBpa mutants were integrated into the *S*. Typhimurium chromosome to maintain native levels of expression (Fig. 1B). Importantly, the resulting strains carrying the *p*Bpa-containing PrgH alleles were fully competent for type III protein secretion (Fig. S3A), indicating that the introduction of the mutations did not affect the assembly nor the function of the T3SS injectisome. UV-mediated crosslinking of PrgH-carrying the different *p*Bpa mutants followed by two-color multiplex Western blot analysis revealed that the replacement of PrgH residues Q42, S43, Q51, I55, or D65 by pBpa resulted in all cases in the detection of a mobility-shifted band of ∼75 kDa (Fig. 1B). The molecular weight of the cross-linked species is consistent with the mass expected for the PrgH-OrgA^3FLAG^ crosslink (44.4 kDa + 25.7 kDa), which was confirmed by Western blotting (Fig. 1B). Quantification of the protein band intensities showed that in some cases (e.g. *PrgH*^*S43pBpa*^) more than half of the detected OrgA protein signal corresponded to crosslinked species, an indication of the robustness of the assay (Fig S4). Mapping the positions that yield the more intense crosslinks onto the solved structure of the cytoplasmic domain of PrgH showed that Q42, S43, and D65 form a cluster located in the FHA inter-strand loops (Fig. 1A and Fig. S2B). The proposed arrangement for the PrgH cytoplasmic domain by the AlphaFold2 model indicates that a lateral surface defined by the β2 and β4 strands of the FHA domain is directly facing OrgA (Fig 1A), raising the possibility that not only the apical loops but also the lateral strands of PrgH contribute to OrgA binding. Such non-canonical binding mode for a FHA domain, in which surfaces other than the loops mediate protein-protein interactions, has been previously described^30,31^. To further probe the accuracy of the PrgH-OrgA organization predicted by the AlphaFold2 model we placed *p*Bpa at the positions of residues C27, E28, and F61 (located in the β2 and β4 strands of PrgH, Fig S2A), which are predicted to be in close proximity to OrgA (Fig. 1A). Introduction of *p*Bpa at any of these positions resulted in a functional T3SS (Fig S3B). Consistent with the predicted model, placement of *p*Bpa in C27 and E28, and to a lesser extent F61, resulted in robust crosslinks with OrgA (Fig. 1C and S4), suggesting that PrgH lateral strands also contribute to the proper docking of the sorting platform into the needle complex.

**Figure 1.**
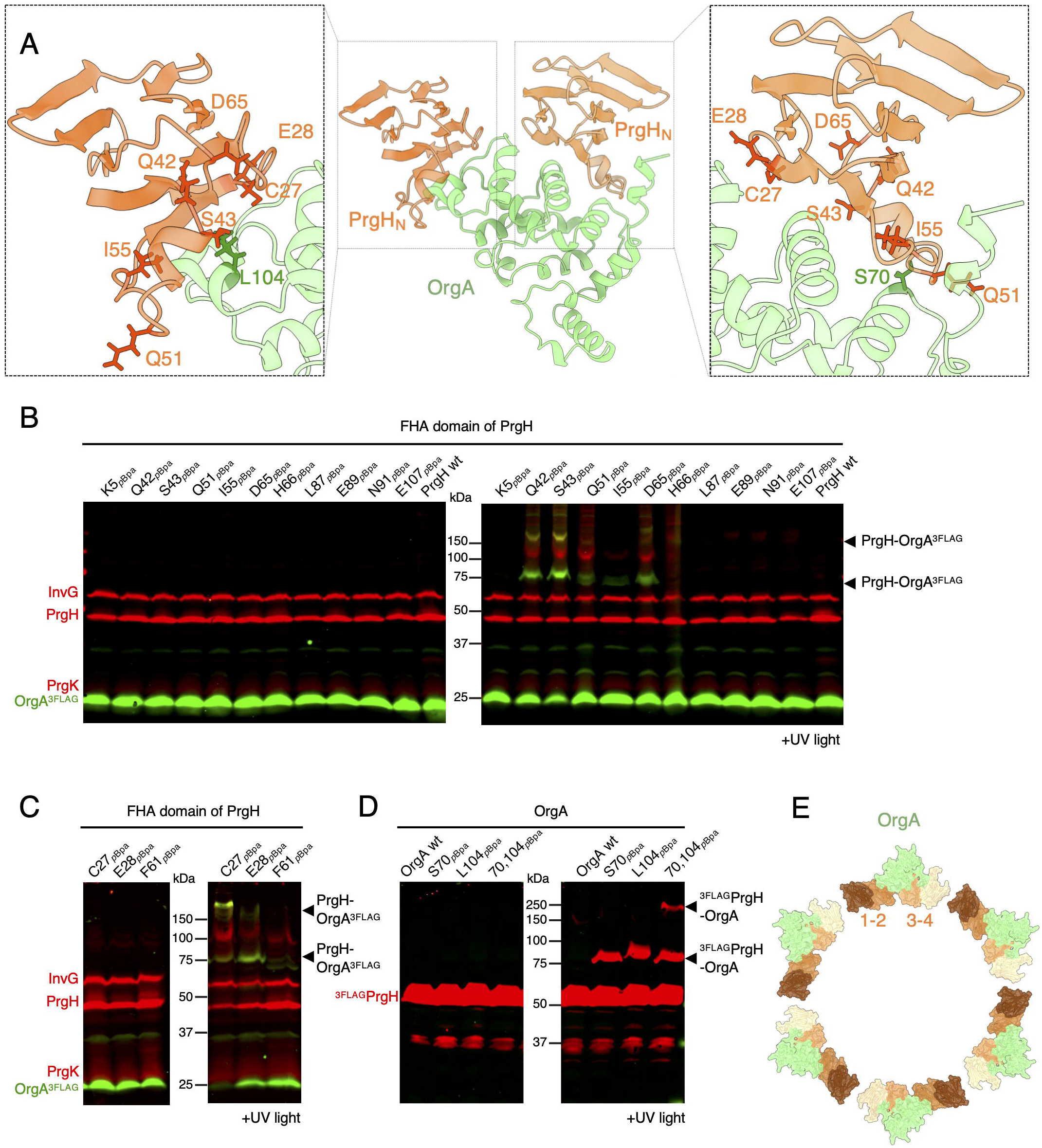
Coupling AlphaFold2 modelling and *in vivo* photocrosslinking to map the needle-complex interface with the sorting platform protein OrgA. (**A**) Structure of the 2:1 PrgH-OrgA complex generated using AlphaFold-Multimer (PrgH in tan, OrgA in green). Insets show the crosslinkable residues in the PrgH-OrgA interfaces mapped onto the structures. (**B** and **C**) Whole cell lysates of *S*. Typhimurium strains expressing OrgA^3FLAG^ and the indicated *p*Bpa-containing PrgH mutants after exposure to UV light or left untreated. Samples were analyzed by Western blot with antibodies to the NC base (red channel, anti-rabbit) or the FLAG epitope (green channel, anti-mouse) in OrgA. (**D**) Whole cell lysates of *S*. Typhimurium strains expressing ^3FLAG^PrgH and the indicated *p*Bpa-containing OrgA variant that have been exposed to UV light or left untreated. Samples were analyzed by Western blot with antibodies to the FLAG epitope. (**E**) Bottom view of the proposed rearrangement of the PrgH_N_ ring upon OrgA docking. Western-blots shown in all panels are representative of three biological replicates.

To further probe the putative OrgA-PrgH binding interface, we introduced *p*Bpa within the region of OrgA that AF2 predicted to interact with PrgH_N_ and carried out the reciprocal cross-linking experiments as described above. We found that placement of *p*Bpa at OrgA residues S70, R100, L104, and L105 resulted in UV-dependent cross-links with mobilities compatible with a PrgH-OrgA^3FLAG^ complex (Fig S5). The polyclonal antibodies raised against PrgH were not sensitive enough to detect PrgH in these crosslinked complexes (Fig S5). However, UV-irradiation of *S*. Typhimurium strain encoding OrgA^S70*p*Bpa^ or OrgA^L104*p*Bpa^ as well as ^3FLAG^PrgH corroborated the identity of the crosslinked adducts (Fig 1D). Interestingly, in the predicted AF2 model, the residues S70 and L104 are located in different OrgA lobes, each one interfacing a distinct PrgH cytoplasmic subunit (Fig 1A). To confirm that, as predicted by the AF2 model, OrgA simultaneously interacts with two different PrgH_N_ subunits, we combined both pBpa mutations (S70*p*Bpa and L104*p*Bpa) into a single OrgA in a strain expressing ^3FLAG^PrgH. Importantly, none of these mutations impair T3SS function (Fig S3C). After UV irradiation, in addition to the ∼75 kDa complex corresponding to the OrgA-^3xFLAG^PrgH crosslink observed in the single *p*Bpa derivatives, the double *p*Bpa-containing OrgA^S70*p*Bpa/L104*p*Bpa^ strain produced an additional band of higher molecular weight containing ^3xFLAG^PrgH indicating that two PrgH subunits simultaneously crosslink to a single OrgA (Fig 1D).

Taken together, these results show that the cytoplasmic domains of two molecules of PrgH directly interact with a single molecule of OrgA through their FHA domains, formally demonstrating that OrgA directly links the sorting platform to the needle complex. Importantly, the proposed organization of the PrgH-OrgA interface provides a mechanistic explanation for the observed reorganization of the cytoplasmic domain of PrgH upon OrgA binding, which results in the formation of 6 discrete patches predicted to contain 4 molecules of PrgH (see above). In fact, higher-order adducts were detected in several of our cross-linked samples (Fig. 1B), suggesting the capture of a PrgH-OrgA complex in a more intricate arrangement, most likely representing the reported rearrangement of the cytoplasmic domain of PrgH^18^. We proposed that the observed arrangement of the cytoplasmic domains of two PrgH molecules each with a distinct orientation and in direct contact with OrgA triggers a conformational change in the immediately adjacent subunits not bound to OrgA rendering them unable to interact with their neighboring subunits on the other side, thus resulting in the formation of the observed 4-subunit patches on the interface with OrgA (Fig 1E and S6).

### PrgH-bound OrgA serves as a platform for the docking of SpaO

We tested whether the scaffold formed by PrgH-bound OrgA could serve as the docking site for SpaO, which is the major component of the *Salmonella* sorting platform pods. The C-terminal half of SpaO contains two domains known as <u>s</u>urface <u>p</u>resentation <u>o</u>f <u>a</u>ntigens domains 1 (SPOA1) and 2 (SPOA2) (Fig S7A) whose structure has been solved^32^. These two C-terminal SPOA domains serve as a protein-interaction hub engaging in multiple interactions with other binding partners including OrgB^32,33^. In contrast, the structural and functional details of the N-terminal half of the SpaO (SpaO_N_) have remained elusive, although a previous mutagenesis study found residues in this region that are critical for its function^33^. We modeled the OrgA-SpaO complex with AlphaFold2, which predicted that the amino-terminal domain of SpaO mediates its association to OrgA (Fig. 2A). Therefore, we introduced *p*Bpa throughout the amino terminal domain of SpaO and probed for potential protein-protein interactions. Most of the introduced mutations did not affect T3SS function (Fig S7B). Of all the positions tested, *p*Bpa incorporation at position 88 of SpaO produced a weak but reproducible crosslink with OrgA indicating a spatial proximity of this SpaO domain to OrgA (Fig. S7C). To narrow down the precise SpaO region that interacts with OrgA, we systematically introduced *p*Bpa at residues surrounding SpaO^E88^ (positions L84, A85, A86, T87, R89, and F91), which did not affect the function of the T3SS (Fig. S8A). We found that, after UV exposure, *p*Bpa placed at SpaO^T87^ yielded a robust mobility-shifted band of ∼60 kDa, which is compatible with a 1:1 complex between OrgA and SpaO (25.7 kDa+35.9 kDa) (Fig. 2B and S9). Consistent with this hypothesis, two-color Western blot showed that this crosslinked adduct contained both OrgA^3FLAG^ and ^2HA^SpaO (Fig. 2B), corroborating that this SpaO region forms its contact surface with OrgA. Consistent with the AF2 predicted model, SpaO T87, which when replaced by *p*Bpa showed robust cross-linking, is predicted to be spatially very close (< 5Å) to OrgA (Fig. 2A). Similarly, the predicted OrgA-SpaO interface also revealed a loop in the N-terminal region of OrgA, which is located in close proximity to the predicted SpaO-interacting region of the amino terminal domain of OrgA (Fig. 2A). Incorporation of *p*Bpa into this loop (OrgA^E38*p*BpA^) did not exert a negative effect on T3SS activity (Fig S8B) and resulted in a UV-dependent crosslinked species of ∼60 kDa, which multiplex Western blotting showed to contain both OrgA^3FLAG^ and ^2HA^SpaO (Fig. 2Cand S9B). These results indicate that the N-terminal regions of OrgA and SpaO interact with each other to assemble the pods of the *Salmonella* sorting platform. Consistent with this notion mutations in residues mapping to this region of OrgA and predicted to be involved in this interaction (i. e. I35K and L52S), significantly impaired T3SS function (Fig S10A), and prevented its crosslinking with SpaO^T87*p*Bpa^ (Fig S10B). Taken together, these data support the notion that PrgH-bound OrgA serves as a platform that engages the N-terminal domain of SpaO.

**Figure 2.**
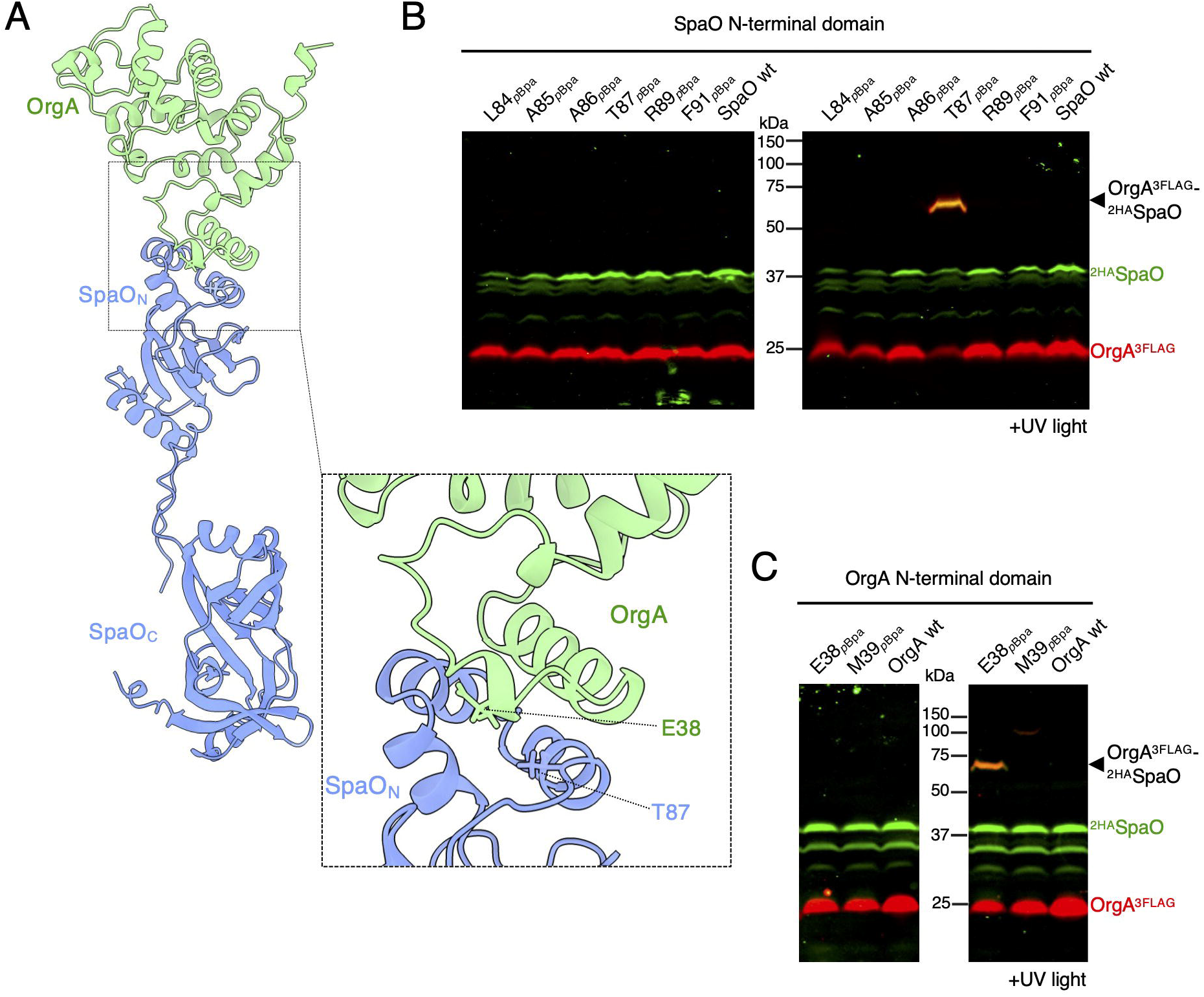
Defining the SpaO-OrgA interface. (**A**) AlphaFold-Multimer model of the OrgA-SpaO complex (OrgA in green, SpaO in blue). Inset shows, the crosslinkable residues between OrgA and SpaO mapped onto the structures. (**B** and **C**) Whole cell lysates of *S*. Typhimurium strains chromosomally expressing OrgA^3FLAG^ (**B**) or ^2HA^SpaO (**C**) and the indicated *p*Bpa-containing ^2HA^SpaO (**B**) or OrgA^3FLAG^ (**C**) variants, that have been exposed to UV light or left untreated. Samples were analyzed by Western blot with antibodies directed to the HA (green channel, anti-mouse) or FLAG (red channel, anti-rabbit) epitopes. Western-blots shown in all panels are representative of three biological replicates.

### Assembly of the sorting platform pods

In contrast to the NC and the export apparatus, the precise mechanisms that lead to the assembly of the sorting platform are currently not understood. The definition of the interacting surfaces between the different components of the sorting platform pods, coupled with the ability to detect specific pairwise interactions between these components *in vivo* through photo-crosslinking (see above), afforded us the opportunity to track the assembly state of the sorting platform in live cells, with high spatial resolution, and in the context of the presence or absence of specific components of the T3SS. To investigate the specific requirements for OrgA recruitment to the needle complex, we introduced the *PrgH*^*S43pBpa*^ allele into *S*. Typhimurium strains carrying deletions in the genes encoding each one of the components of the sorting platform (Fig. 3A). As shown above, PrgH^S43*p*Bpa^ can be efficiently crosslinked to OrgA upon exposure to UV light. Therefore, this mutant can serve as a reporter for OrgA recruitment to the needle complex *in vivo*, thus allowing us to examine the specific requirement of each individual components of the sorting platform for the OrgA recruitment process. We found that the recruitment of OrgA onto PrgH occurs efficiently in the absence of any other components of the sorting platform (i.e. OrgB, SpaO, InvC, or InvI) (Fig. 3B), although as previously reported^34^, its stability was somewhat compromised in the absence of SpaO. Taken together these results indicate that OrgA has intrinsic affinity for PrgH that is not dependent on the presence of other components of the sorting platform.

**Figure 3.**
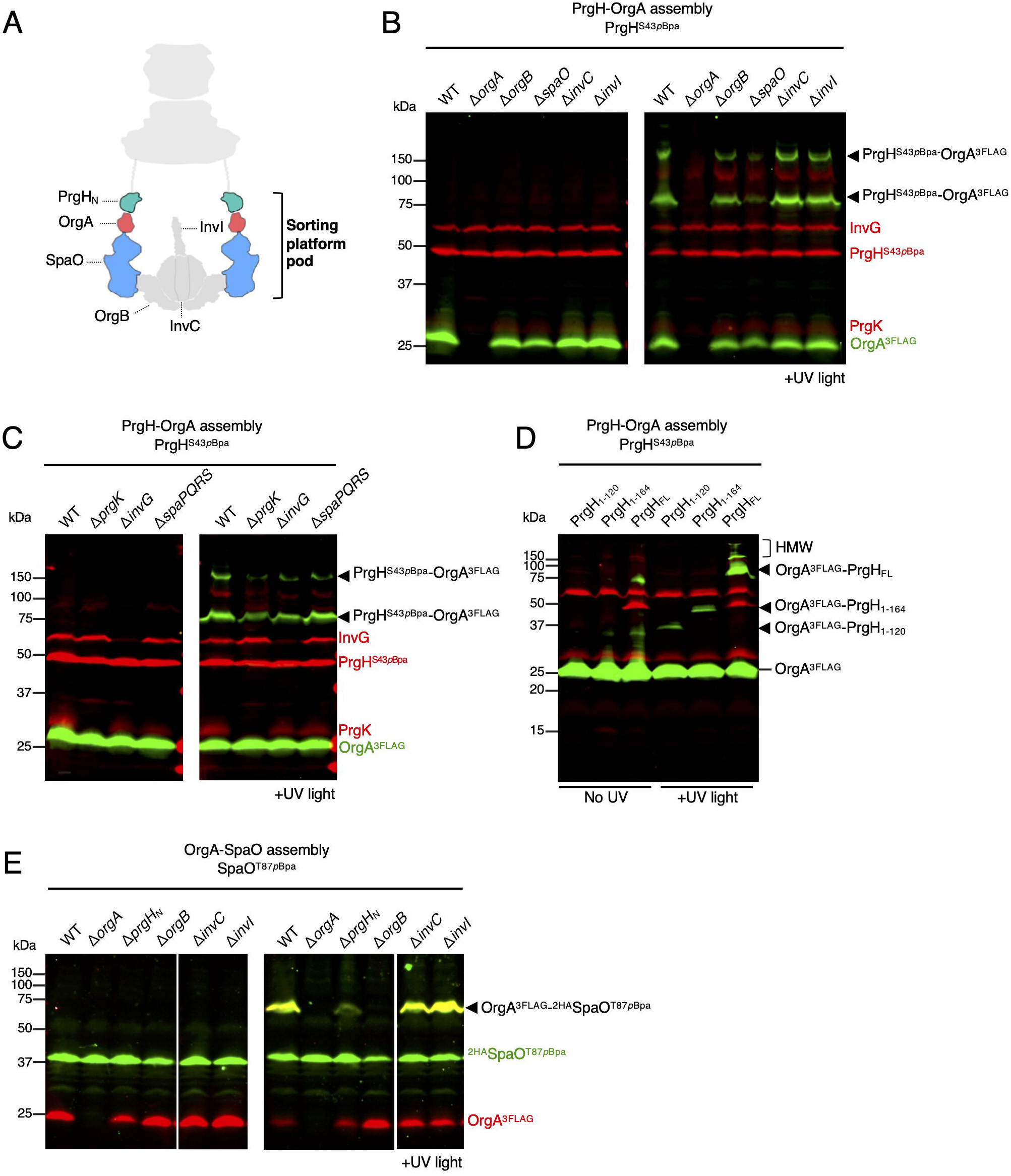
Dissecting the assembly pathway of the sorting platform pods. (**A**) Schematic diagram of the *Salmonella* T3SS sorting platform pods. (**B** and **C**) OrgA binds to PrgH independently of the sorting platform, needle-complex, or export apparatus subunits. Whole cell lysates of *S*. Typhimurium strains expressing OrgA^3FLAG^ and PrgH^S43*p*Bpa^ in combination with the indicated deletions of genes encoding sorting platform (**B**) or needle-complex and export apparatus components (**C**), were analyzed (before and after UV light exposure, as indicated) by Western blot with antibodies directed to the NC base (red channel, anti-rabbit) or the FLAG epitope (green channel, anti-mouse). (**D**) Whole-cell lysates of *S*. Typhimurium encoding full-length (PrgH_FL_) or truncated forms of PrgH^S43*p*Bpa^ allele were analyzed (before and after UV light exposure, as indicated) by Western blot with antibodies directed to the NC base (red channel, anti-rabbit) or the FLAG epitope (green channel, anti-mouse). HMW, high molecular weight adducts. (**E**) Assembly profile of the OrgA-SpaO complex. The analysis of whole cell lysates was conducted as in (**B**) but employing the reporter strain ^2HA^SpaO^T87pBpa^ and using antibodies directed against the HA (green channel) or FLAG (red channel) epitopes. Western-blots shown in all panels are representative of three biological replicates.

It is generally assumed that the full completion of the assembly of the of needle-complex base substructure must precede the recruitment and docking of the sorting platform components^4^. However, we found efficient crosslinking of OrgA to PrgH^S43*p*Bpa^ even in the absence of other components of the needle complex base (i. e. PrgK or InvG), or the export apparatus (i. e. SpaP/Q/R/S) (Fig. 3C). PrgH is capable of forming a multimeric structure by itself. Therefore, we tested whether the docking of OrgA onto PrgH requires its oligomerization. To this aim, we generated truncated versions of PrgH^S43*p*Bpa^ resulting in the production of only the cytoplasmic FHA domain (PrgH_1-120_, i. e. the OrgA binding domain) or the cytoplasmic FHA domain plus the transmembrane helix (PrgH_1-164_) by introducing TAA premature termination codons into the PrgH^S43*p*Bpa^ chromosomal allele (Fig S11A). As expected, the truncated versions of PrgH^S43*p*Bpa^ lacking the entire periplasmic domain that forms the inner membrane ring upon oligomerization, resulted in a non-functional type III secretion system (Fig S11B). Exposure of these strains to UV light, followed by immunodetection of crosslinked species showed that OrgA can bind the FHA domain of either of these PrgH constructs (Fig. 3D). These results indicate that OrgA can bind PrgH even in the absence of its oligomerization. In contrast, the formation of higher-order structures between PrgH and OrgA were only observed in the presence of full-length PrgH (Fig 3D), suggesting that the complex rearrangements in PrgH_N_ triggered by the sorting platform incorporation take place only in the context of a fully assembled PrgH ring. Taken together, these results indicate that OrgA can engage PrgH even in the absence of either the sorting platform or the needle-complex components.

Next, we evaluated the requirements for the formation of the OrgA-SpaO complex. To this aim, we made use of SpaO^T87*p*Bpa^, which exhibited robust cross-linking to OrgA. We introduced this allele into *S*. Typhimurium strains carrying deletions in each one of the genes encoding the components of the sorting platform, as well as into a strain carrying a deletion in the cytoplasmic domain of PrgH, which serves as binding site for OrgA. We then examined the presence of the OrgA-SpaO crosslink species after exposure to UV light of the resulting strains. We found that in the absence of the cytoplasmic domain of PrgH, the OrgA-SpaO complex could still be detected although at lower levels than in the presence of wild type PrgH (Fig. 3E). Although some of the reduced ability of OrgA to engage SpaO in the absence of the cytoplasmic domain of PrgH may be explained by its reduced stability in this strain background, it is clear that formation of the OrgA-SpaO complex is compromised in the absence of PrgH. This is in contrast with the formation of the OrgA-PrgH complex, which was minimally affected by the absence of SpaO (Fig. 3B). The OrgA-SpaO complex was readily detected in the absence of InvI or InvC, indicating that a functional type III secretion is not required for the stable association of these sorting platform components. In contrast, this complex could not be detected in the absence of OrgB, even though the stability of SpaO and OrgA was unaltered in this strain (Fig. 3E). Introduction of a plasmid expressing wild type OrgB into the *ΔorgB* strain restored the formation of the SpaO-OrgA complex (Fig S12A and S12B). Furthermore, the OrgA-SpaO complex was also undetectable in a strain expressing a mutant of SpaO (SpaO^G289D^) (Fig S12C and S12D), which is unable to interact with OrgB^34^. These observations indicate that SpaO must form a complex with OrgB to become competent for binding to OrgA. This is surprising since OrgB interacts with the carboxy-terminal domain of SpaO, away from its OrgA interacting region. Therefore, binding to OrgB likely triggers a conformational change in SpaO, which is necessary to render it competent for OrgA binding. Taken together, these results showed a highly coordinated process that leads to the assembly of the sorting platform pods, initiated by the binding OrgA to the cytoplasmic domain of PrgH, and followed by the recruitment of OrgB-bound SpaO to the PrgH-OrgA complex.

### Topological organization of the *Salmonella* sorting platform ‘cradle’

The six-pod cage-like structure formed by OrgA and SpaO is capped at its most distal end by a six-spoke wheel-like structure formed by an array of OrgB dimers^18^. It is hypothesized that this scaffold arrangement provides a ‘cradle’ for the ATPase InvC just beneath the entry to a multi-ring nonameric assembly formed by the cytoplasmic domain of InvA. At this site, InvC can couple its unfoldase and chaperone-stripping activities to the protein export process^35^. The X-ray structure of the SPOA domains of SpaO in complex with an amino terminal fragment of OrgB showed that the disordered N-terminal end of OrgB binds to both the SPOA1 and SPOA2 domains of the C-terminus of SpaO^32^. On the other hand, the ATPase is proposed to be anchored to the sorting platform pods presumably by interactions with the C-terminal domain of OrgB^36^. However, no *in-vivo* evidence has been reported to confirm this proposed arrangement. To investigate the topological organization of the sorting platform cradle *in vivo*, we incorporated *p*Bpa into the N-terminal domain of OrgB at positions K20 and R21, which the crystal structure of the SpaO-OrgB complex predicts to be in close proximity to SpaO (Fig. 4A). A functional assessment showed that both mutant strains were fully competent for type III secretion indicating that introduction of *p*Bpa at these OrgB locations did not alter its function (Fig S13A). Exposure of a *S*. Typhimurium expressing OrgB^R21*p*Bpa^ to UV light resulted in the appearance of a crosslinked product of ∼50 kDa compatible with an OrgB dimer (Fig. 4B). In contrast, exposure to UV light of a strain expressing OrgB^K20*p*Bpa^ resulted in the detection of a crosslinked band with a larger molecular size, which upon multiplex immunoblotting was shown to contain both OrgB and SpaO (Fig. 4B). The apparent molecular weight of the cross-linked adduct (∼100 kDa) is consistent with a complex formed by 2 molecules of OrgB and 1 molecule of SpaO (see below).

**Figure 4.**
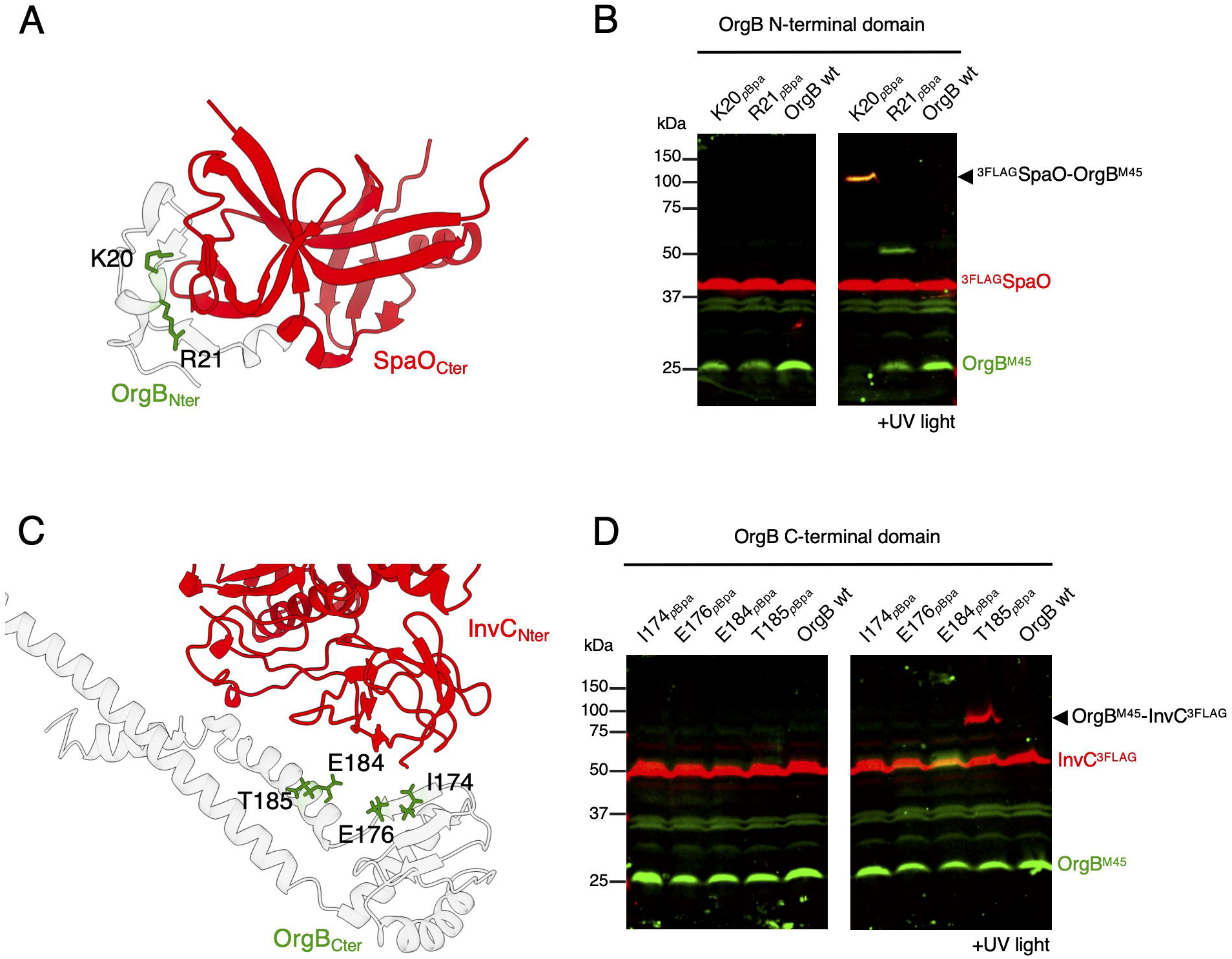
Systematic OrgB photocrosslinking analyses identifies the sorting platform cradle interfaces. (**A and B**) Identification of crosslinkable amino acids in the OrgB-SpaO interface. Crystal structure of SpaO (red) in complex with the N-terminal region of OrgB_1-30_ (gray) (PDB ID code 4YX7) showing (in green) the OrgB residues (K20 and R21) selected for *p*Bpa incorporation (**A**). Whole cell lysates of *S*. Typhimurium strains expressing ^3FLAG^SpaO and the indicated OrgB^M45^ *p*Bpa variants after exposure to UV light (or left untreated) were analyzed by Western blot with antibodies directed against the FLAG (red channel, anti-rabbit) or M45 (green channel, anti-mouse) epitopes (**B**). (**C** and **D**) Structural modeling and site directed photocrosslinking define OrgB-InvC interface. Enlarged view of the modeled InvC (red)-OrgB (grey) interface (**C**). The OrgB residues chosen for *p*Bpa incorporation are shown in green (**C**). Whole cell lysates of *S*. Typhimurium strains expressing InvC^3FLAG^ and the indicated OrgB^M45^ *p*Bpa variant, exposed to UV light or left untreated, were analyzed by immunoblot using antibodies directed to the FLAG (red channel, anti-rabbit) or M45 (green channel, anti-mouse) epitopes. The crosslinks between InvC^3FLAG^ and OrgB^M45^ are indicated. Western-blots shown in all panels are representative of three biological replicates.

We then investigated the structural organization of the OrgB-InvC complex. While the structure of the InvC monomer has been partially solved ^37^, the structural details of the InvC-OrgB complex are not available. However, the flagellar ATPase FliI has been co-crystallized in complex with a dimer of FliH^38^, the flagellar OrgB homolog. Therefore, using this structure as a template (PDB ID code 5B0O), we modeled the structure the OrgB-InvC complex and identified 4 residues at the C-terminal end of OrgB (I174, E176, E184, and T185), which, according to the model, are predicted to be in close proximity to InvC (Fig. 4C). We then replaced those residues with *p*Bpa, and introduced the mutated alleles into the *S*. Typhimurium chromosome. The resulting strains retained type III secretion function in a manner indistinguishable from wild type (Fig S13B). Exposure of the *S*. Typhimurium strain expressing the OrgB^T185*p*Bpa^ allele to UV light resulted in the formation of a crosslinked product of ∼75 kDa consistent with a crosslinked 1:1 complex between OrgB^M45^ and InvC^3FLAG^ (28.7 kDa+ 50.5 kDa), an arrangement that was confirmed by multiplex immunoblotting (Fig. 4D). Taken together, these results define the molecular interfaces between OrgB and its interacting partner SpaO, as well as the docking site for the ATPase InvC on the sorting platform cradle.

### Assembly of the *Salmonella* sorting platform ‘cradle’

To obtain insight into the assembly of the sorting platform cradle and the recruitment of InvC to the sorting platform, we investigated the formation of the OrgB-SpaO or OrgB-InvC complexes in the absence of specific components of the sorting platform or the cytoplasmic domain of PrgH. To monitor the formation of the OrgB-SpaO and OrgB-InvC complexes in the mutant strains, we introduced the OrgB^K20*p*Bpa^ or the OrgB^T185*p*Bpa^ alleles (see above), respectively, into the different mutant strains. We found that formation of the OrgB-SpaO complex was readily detected in the absence of the cytoplasmic domain of PrgH, indicating that formation of this sub-complex can occur in the absence of fully assembled sorting platform (Fig. 5A and 5B). Consistent with this notion, assembly of the OrgB-SpaO complex was also detected in the absence of OrgA, which links the sorting platform pods to the needle complex (Fig. 5B). We noticed, however, that the deletion of *orgA* resulted in a marked reduction in the levels of OrgB. The *orgA* gene is located upstream of *orgB*, sharing an overlap of 44 bp in their coding sequences (Fig S14A). Such genomic arrangement in combination with the observed reduction in the levels of OrgB suggested that *orgB* may be translationally coupled to *orgA*, or that promoter sequences located within *orgA*^39^, may influence its expression. Therefore, to examine the assembly state of the OrgB-SpaO in the absence of *orgA* under conditions that would eliminate any potential confounding transcriptional or translational effects derived from the genomic organization, we constructed strain in which *orgA* was replaced by a synthetic RBS that ensures efficient *orgB* translation (Fig S14). We found that in this strain expression levels of OrgB were similar to those of wild type, indicating that, as hypothesized, deletion of *orgA* altered the transcription and/or translation of *orgB* (Fig. 5C). Importantly, however, formation of the OrgB^K20*p*Bpa^-SpaO complex in this background strain was similar to what was observed in the wild type strain (Fig. 5C), corroborating that the absence of OrgA does not influence the formation of the SpaO-OrgB complex. Similarly, absence of InvC or InvI had no impact on the formation of the OrgB-SpaO complex (Fig 5B). These results indicate that formation of the SpaO-OrgB subcomplex does not require any other sorting platform component, suggesting that formation of this complex must precede the initiation of the sorting platform assembly process. On the other hand, formation of the OrgB-InvC complex could not be detected in the absence of any of the sorting platform pod components or the cytoplasmic domain of PrgH, indicating that a fully assembled sorting platform is required to recruit InvC (Fig. 5D). Of note, recruitment of InvC to the sorting platform did not require a functional type III secretion system since it efficiently occurred in the absence of InvI (Fig. 5D), an essential component of the T3SS that links the ATPase to the export gate.

**Figure 5.**
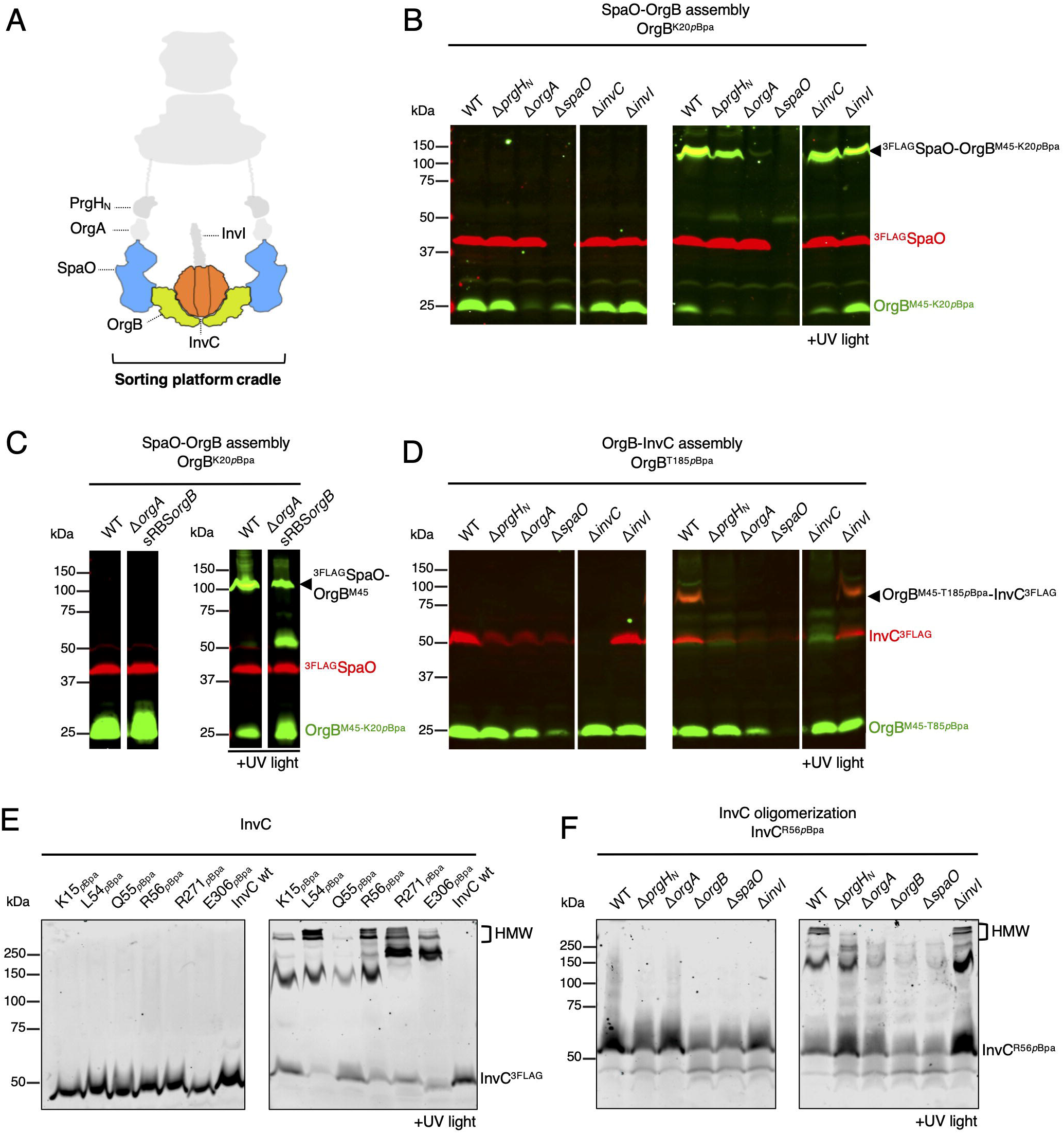
Monitoring the assembly of the sorting platform cradle. (**A**) Schematic diagram of the *Salmonella* T3SS sorting platform cradle. (**B** and **C**) Formation of the SpaO-OrgB complex does not require an assembled sorting platform. Whole-cell lysates of *S*. Typhimurium wild-type or the indicated isogenic mutant strain expressing OrgB^K20*p*Bpa^ and ^3FLAG^SpaO, exposed to UV light or left untreated, were analyzed by Western blot with antibodies directed to the M45 (green channel, anti-mouse) or FLAG (red channel, anti-rabbit) epitopes. (**D**) InvC docking onto the OrgB cradle requires a fully assembled sorting platform. Whole cell lysates of *S*. Typhimurium wild-type or the indicated isogenic mutant strain expressing OrgB^T185*p*Bpa^ and InvC^3FLAG^ analyzed as indicated in (**B**) and (**C**). (**E**) Survey for crosslinkable sites in the InvC hexameric interfaces. Whole cell lysates of *S*. Typhimurium strains expressing the indicated *p*Bpa-containing InvC^3FLAG^ variant, exposed to UV light or left untreated, were analyzed by immunoblot with antibodies directed to the FLAG epitope. (**F**) Monitoring the oligomerization of the InvC ATPase. Whole-cell lysates of *S*. Typhimurium strains expressing the InvC^R56*p*Bpa^ allele in combination with the indicated deletion mutants, exposed to UV light or left untreated, were analyzed by western-blot with antibodies directed to the FLAG epitope. HMW, high molecular weight adducts. Western-blots shown in all panels are representative of three biological replicates.

Oligomerization of T3SS ATPases is needed to fully exert its catalytic activity and hence for proper type III protein export^40-42^. To investigate whether oligomerization of InvC requires its recruitment to a fully assembled sorting platform we sought to introduce *p*Bpa at positions located within the inter-subunit interface so that they could report inter-subunit interactions *in vivo*. Although the hexameric structure of InvC is not available, the cryo-EM map of the ortholog ATPase EscN from enteropathogenic *E. coli* was recently obtained^43^. Thus, we modelled the InvC hexamer using the EscN cryo-EM structure as a template and guided by this model, we introduced pBpa at 6 positions (K15, L54, Q55, R56, R271 and E306) that the modelled complex predicted to be located in the inter-subunit interface (Fig S15A). All mutants but one (InvC^E306pBpa^) were stably expressed at levels comparable to wild type (Fig 5E). Furthermore, all mutant strains were fully competent for type III secretion except for InvC^E306*p*Bpa^ that showed reduced secretion, and InvC^L54pBpa^, which completely lacked type III protein secretion activity (Fig S15B). Exposure of the *p*Bpa-containing InvC derivatives to UV light resulted in a ladder of crosslinks of high-molecular weight that we interpret to represent oligomeric forms of InvC (Fig. 5E). To evaluate the requirements for InvC assembly into an oligomeric structure we used a *S*. Typhimurium strain expressing InvC^R56pBpa^, which showed the most robust crosslinks due to InvC oligomerization. We then introduced in this reporter strain deletion mutations in genes encoding sorting platform components and examined InvC oligomerization. We found that removal of any of the sorting platform components except for InvI abolished the appearance of high-molecular weight species of InvC^R56*p*Bpa^ (Fig. 5F), indicating that full oligomerization of InvC occurs only when the sorting platform scaffold is fully formed. Furthermore, the observation that the InvC^R56*p*Bpa^ crosslinks could be detected in the absence of InvI indicates that InvC oligomerization does not require an active type III secretion machine.

### Architecture of the core sorting platform components

The protein-protein interactions between PrgH-OrgA and between OrgA-SpaO captured in our crosslinking studies support the notion that OrgA acts as a structural linker between the needle complex and the sorting platform. Previous studies have reported an estimated relative stoichiometry of 6 OrgA copies per injectisome^44,45^. Since there are six pods in a sorting platform, the observed stoichiometry suggests that a single molecule of OrgA would interact with both PrgH and SpaO. However, no direct proof for such arrangement has been reported. To test this hypothesis, we constructed a *S*. Typhimurium strain that expresses both the *prgH* (*prgH*^*S43pBpa*^) and *spaO* (*spaO*^*T87pBpa*^) alleles carrying *p*Bpa substitutions at positions that resulted in robust crosslinks to OrgA (see above). If the same molecule of OrgA links PrgH and SpaO, exposure of this strain to UV light should result in a crosslink that contains the three proteins. As shown in Fig. 6A and 6B, we found that exposure of this strain to UV light resulted in the formation of a crosslinked adduct of higher molecular weight than the crosslinks observed with each one of the single *p*Bpa derivative strains (Fig 6A and B). This observation indicates that a single OrgA molecule links SpaO to the cytoplasmic domain of PrgH thus bridging the individual pods of the sorting platform to the needle complex.

**Fig 6.**
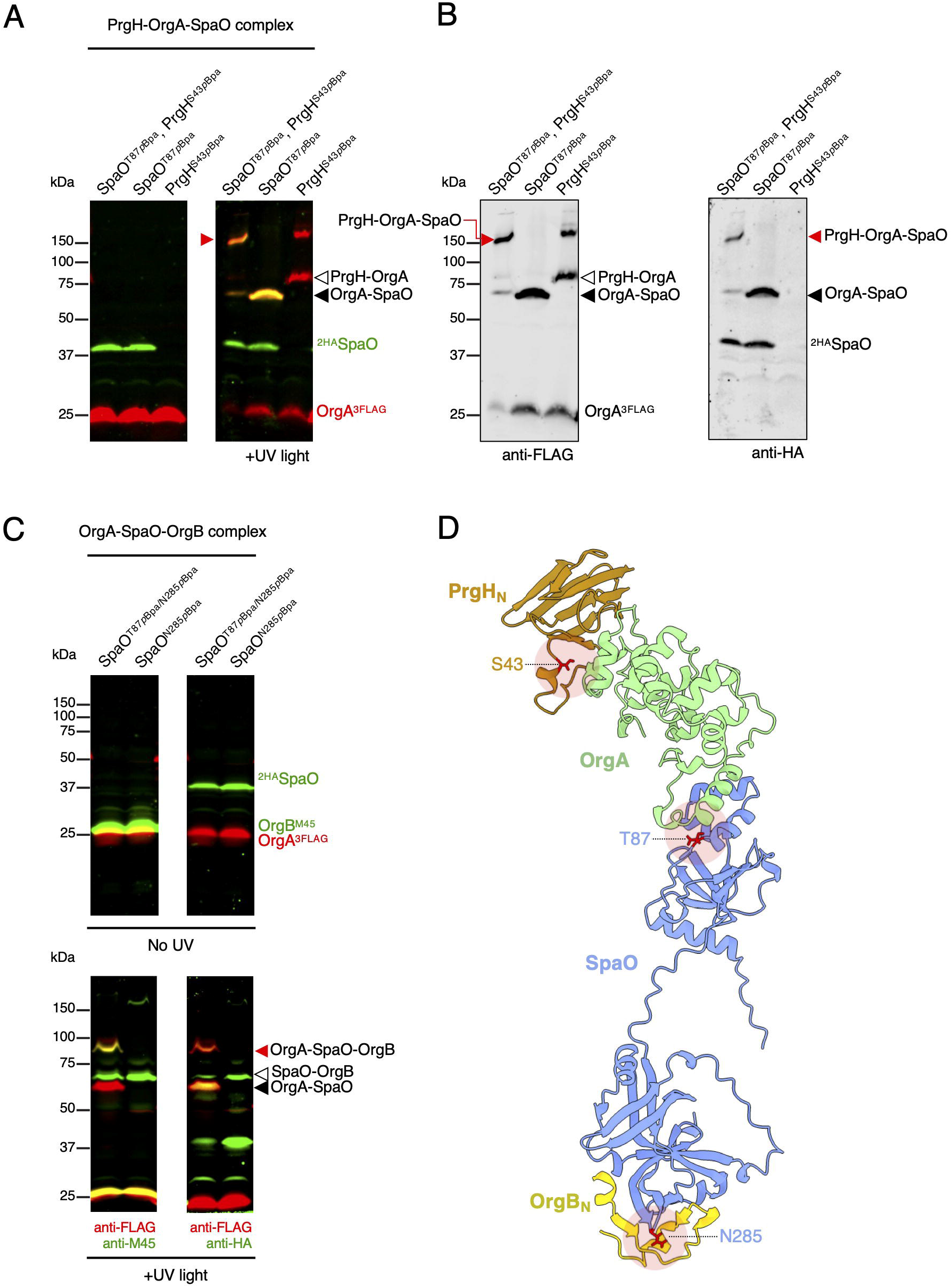
Architecture of the sorting platform. (**A** and **B**) The OrgA protein tethers the sorting platform to the needle-complex. Whole cell lysates of *S*. Typhimurium strains expressing OrgA^3FLAG^ in conjunction with ^2HA^SpaO^T87pBpa^, PrgH^S43pBpa^, or both, as indicated, either exposed to UV light or left untreated. Lysates were analyzed by Western blot with antibodies directed to the HA (green channel, anti-mouse) or FLAG (red channel, anti-rabbit) epitopes (**A**). The individual acquisition channels for anti-FLAG (Odyssey scan at channel 700 nm, red in panel **A**) and anti-HA (Odyssey scan at channel 800 nm, green in panel **A**) in the UV-exposed samples are shown (**B**). The position of the tripartite and bipartite complexes containing the indicated proteins are noted. (**C**) A single SpaO subunit bridges OrgA and OrgB. Whole cell lysates of *S*. Typhimurium strains expressing OrgA^3FLAG^, OrgB^M45^ and either ^2HA^SpaO^T87*p*Bpa/N285*p*Bpa^ or ^2HA^SpaO^N285*p*Bpa^ after exposure to UV light or left untreated. Cell lysates were analyzed by Western blot with antibodies directed to the FLAG and M45 epitopes (left panels), or to the FLAG and HA epitopes (right panels). The position of the tripartite and bipartite complexes containing the indicated proteins are noted. (**D**) Ribbon diagram of the model for the PrgH-OrgA-SpaO-OrgB complex (PrgH in tan, OrgA in green, SpaO in blue, OrgB in yellow). The PrgH-OrgA-SpaO complex was predicted using AlphaFold2 and the OrgB_N_ fragment was placed by aligning the structure of the SpaO-OrgB complex onto the AF2 model. The crosslinkable residues employed in the experiments shown here are highlighted in red.

Likewise, the precise composition of the structural core forming the sorting platform pods is uncertain. It has been estimated that there are 24 copies of SpaO per needle complex^44,45^, a relative stoichiometry that would result in 4 SpaO subunits per pod. However, the direct cryo-ET visualization of the sorting platform^18^ lacks the resolution required to resolve the atomic spatial arrangement of its building blocks and confirm this proposed arrangement. In addition, based mainly on protein modeling and fitting into the cryo-ET density map, a previous report suggested that each pod is composed of two copies of SpaO per pod, one interfacing with OrgA and the other with OrgB. In this model, each copy of SpaO would necessarily exhibit a different conformation ^46^. However, our observation that the interaction of SpaO with OrgA requires binding of the former to OrgB (Fig. 3E) is at odds with this proposed model. To clarify this issue, we constructed a *S*. Typhimurium strain encoding a 2xHA epitope tagged SpaO (^2xHA^SpaO^T87*p*Bpa/N285*p*Bpa^) containing two pBpa residues at positions that can crosslink with epitope tagged OrgA^3FLAG^ (SpaO^T87*p*BpA^) and OrgB^M45^ (SpaO^N285*p*BpA^). As the discrete N- and C-terminal domains of SpaO interact with OrgA and OrgB, respectively, we reasoned that this strain should be able to inform whether a single SpaO molecule can link the OrgA docking platform to the OrgB cradle. As expected, upon exposure of this mutant strain to UV light and multiplex Western blot analysis, we detected bands corresponding to the SpaO-OrgA and SpaO-OrgB pair-wise crosslinked complexes (Fig. 6C). Importantly, however, we also detected a third UV-dependent adduct of higher molecular weight that contained ^2xHA^SpaO^T87*p*Bpa/N285*p*Bpa^ as well as OrgA^3xFLAG^ and OrgB^M45^. The molecular weight of this adduct, 90 kDa, is consistent with the expected size of the OrgA^3xFLAG^/^2xHA^SpaO/OrgB^M45^ trimeric complex (25.7+35.9+28.7=∼90kDA kDa) (Fig. 6C). Overall, this result indicates that one molecule of SpaO per pod is able to simultaneously link the OrgA docking platform with the OrgB cradle suggesting a topological arrangement for the sorting platform pods (Fig 6D), which differs from a previously proposed model^47^.

## Discussion

Type III secretion systems are among the most complex supramolecular assemblies known to date in bacteria. One of its components, the cytoplasmic sorting platform, plays a central role in the selection and initiation of the various type III secretion clients into the secretion pathway^15^. Although cryo-ET analysis has provided a low-resolution blueprint of the sorting platform organization^18^, the inability to isolate this substructure in a manner amenable to single particle cryo-EM or crystallography studies has prevented its visualization at atomic resolution. Consequently, the lack of high-resolution structural information has impeded studies of its assembly pathway. Here, we developed a series of molecular tools that enable us to study *in vivo* the architecture and assembly of this critical cytoplasmic module of the type III protein secretion machine.

High-resolution fluorescence microscopy studies have provided an accurate measure of the stoichiometry of the different components of the sorting platform injectisome^44,45^. Thus, for the *S*. Typhimurium SPI-1 T3SS sorting platform, it has been estimated that each one of the six pods that make up the sorting platform is composed of one molecule of OrgA, four of SpaO, and two of OrgB. In turn, each one of the pods is proposed to dock onto a platform made up of the cytoplasmic domains of four molecules of the needle complex inner ring protein PrgH. However, how the different structural components organize to produce the final structure has been unclear. *In situ* cryo-ET studies have suggested that in the *S*. Typhimiurium SPI-1 T3SS, OrgA serves as a link between each pod and the needle complex by directly interacting with the cytoplasmic domain of PrgH and the main pod component SpaO^18^. However, no direct evidence for such arrangement has been reported. Using structural modeling with the deep learning algorithm AlphaFold2 in combination with *in vivo* photocrosslinking we were able to establish that OrgA links each one of the pods with a docking platform made by the cytoplasmic domain of two molecules of the needle complex protein PrgH, which are arranged in a different orientation. We hypothesize that the reorientation of the adjacent PrgH molecules triggered by OrgA binding ultimately leads to the reorganization of the PrgH 24-mer ring into the observed six patches, thus adapting its symmetry to dock to the six-pod sorting platform (Fig. 1E and fig. S6). In our model, PrgH subunits adjacent to the OrgA-bound PrgH molecules would be able to join the complex by lateral interactions. However, those interactions would not be possible with subunits on the adjacent patch, which would exhibit a different orientation (Fig. 1A and 1E). This cascade of events would lead to the formation of the 4-subunit patches that are observed in the cryo-ET structure^18^. Our data also indicate that both PrgH molecules engage OrgA with their forkhead-associated (FHA) domain through non-canonical interactions for this class of domains (Fig. 1A). Indeed, FHA domains are known to engage in protein-protein interactions in a phospho-threonine-specific manner^28^. although PrgH lacks key residues responsible for phosphopeptide recognition. We also mapped the OrgA-binding domain to the N-terminal portion of SpaO, a hitherto uncharacterized region of the major component of the sorting platform. Interestingly, a DALI search^48^ with the AlphaFold model revealed structural similarities between the N-terminal domain of SpaO and the middle domain of FliM (Fig. S16). This domain of FliM has been shown to contact FliG, the flagellar component that attaches the C-ring to the MS ring^49^. These observations indicate a conservation of the interaction mode of these distantly related structural elements even across functionally distinct machines.

While *ex-vivo* interactions between PrgH-OrgA, OrgA-SpaO, and SpaO-OrgB have been reported in multiple T3SS-carrying organism^33,50-52^, how these interactions lead to the formation of a sorting platform pod *in vivo* has not been determined. This is critical given the different stoichiometries of these components in the fully assembled structure. Using *in-vivo* crosslinking we were able to determine that the same SpaO molecule is able to interact through its amino terminal domain with OrgA on its membrane proximal side, and through its carboxy terminal domain with OrgB, linking the pod and cradle substructures. Given the existence of an excess of SpaO molecules over other components of the pod substructure, our observations have important implications regarding the potential arrangement of this core component of the sorting platform, and the proposed dynamic behavior of SpaO based on observations on its homologs in other bacterial species. Based on FRAP studies of YscQ, the SpaO homolog in Yersinia spp., it has been proposed that the sorting platform undergoes rapid cycles of assembly and disassembly, a process thought to be critical for its function^53^. However, the rather complex nature of the interactions among the components of the sorting platform that we have mapped in these studies would make such a process challenging. Our observation that the same SpaO molecule establishes contact with both, OrgA and OrgB to form a sorting platform pod coupled to the existence of an excess of SpaO molecules over both OrgA and OrgB (or its homologs in other systems), suggest the possibility that only a subset of SpaO molecules may engage in this dynamic behavior. In this model, only the SpaO molecules not directly in contact with OrgA and OrgB and therefore not engaged in making up the core pod structure, would be subject to cycles of association and disassociation from the sorting platform. In support of this notion, it should be noted that a dynamic behavior has not been demonstrated for sorting platform components other than a homolog of SpaO in *Yersinia* spp^53^. Furthermore, the cryo ET densities within the sorting platform pods presumably associated with SpaO are not sufficient to accommodate four molecules of this component^18^. More experiments will be required to clarify the important issues raised by our studies.

Our studies have also provided a framework for the understanding of the process that leads to a fully assembled sorting platform (Fig. S17). Our results suggest that one of the earliest steps in this process is the association of OrgA with the cytoplasmic domain of the needle complex component PrgH. We found that this association does not require a fully assembled needle complex nor the oligomerization of PrgH. It is therefore possible that assembly of the sorting platform may be concomitant to the assembly of the needle complex base. The PrgH-bound OrgA serves as a platform where the SpaO-OrgB complex can then dock. We find that SpaO cannot bind OrgA unless it is bound to OrgB. This suggests that OrgB binding to SpaO may trigger a conformational change at the amino terminus of SpaO rendering it competent for docking onto the OrgA-PrgH platform. Assembly of the pods must then lead to the assembly of the cradle structure that caps the sorting platform on its cytoplasmic side and that is formed in its entirety by OrgB. The cradle substructure serves as docking site for the hexameric ATPase InvC, which is essential for the initiation of substrates into the secretion pathway. Previous studies have reported that FliI and FliH, the flagellar homologous to InvC and OrgB, respectively, are able to form stable complexes when co-overproduced ^54^. Similarly, purified ATPase FliI tends to oligomerize *in vitro* when incubated with ATP^41^. These observations led to the assumption that the OrgB-InvC subcomplex (or that of its homologs) is pre-assembled in the cytoplasm prior to the formation of the sorting platform scaffold. On the contrary, our data suggest that, under physiological conditions, both InvC assembly onto the C-terminal domain of OrgB and its oligomerization is promoted only by a fully assembled sorting platform scaffold. Such assembly order is in agreement with cryo-ET data, showing the presence of fully assembled sorting platform pods and cradle in the absence of InvC ^18^.

In summary, our findings have provided novel insight into the molecular organization of the sorting platform with implications for its function, as well as a blueprint for the sorting platform assembly pathway. Importantly, the molecular interfaces defined by these studies constitute unique targets for the potential development of novel anti-infective strategies that specifically disable the formation of this structure, which is critical for the virulence of many pathogenic bacteria.

## Methods

### Bacterial strains, plasmids and growth conditions

All strains used in this work are derivatives of *Salmonella enterica* Typhimurium SL1344^55^ and are listed in Table S2. Strains were routinely cultured in lysogeny broth (LB) at 37°C. When needed, bacterial cultures were supplemented with streptomycin (100 μg/mL), tetracycline (10 μg/mL), ampicillin (100 μg/mL), or chloramphenicol (10 μg/mL). Genetic modifications were introduced in the *S*. Typhimurium chromosome by allelic exchange using the R6K-derived suicide vector pSB890 and *E. coli* β-2163 Δnic35 as the conjugative donor strain as previously described^56^. Molecular cloning was performed using the standard Gibson assembly protocol^57^. To prevent polar effects on downstream genes within the same operon, a *prgK* knock-out strain was constructed in which PrgK production was abolished by chromosomally mutating the ATG start codon of *prgK* to ATA. The presence of the mutant allele was further confirmed by sequencing and abolishment of translation was verified by western-blot analysis and functional assays. To circumvent the low expression levels of *orgB* observed in the Δ*orgA* clean deletion mutant, a synthetic ribosome binding site was designed using the RBS Calculator tool^58^ and placed upstream of the *orgB* gene, instead of the *orgA* gene, to maximize *orgB* translation rate (Fig. S14).

### Cell lines

Henle-407 human epithelial cells were cultured in Dulbecco’s modified Eagle medium (DMEM, Gibco) supplemented with 10% bovine calf serum (BCS) at 37°C under a 5% CO_2_ atmosphere.

### Type III protein secretion assay

Overnight cultures of the different *S*. Typhimurium strains were grown at 37°C in LB supplemented with the proper antibiotics. The ON cultures were used to inoculate (1:20) 10 ml of pre-warmed LB supplemented with 0.3 M NaCl and subcultures were grown under low aeration until an OD_600_ ∼ 0.8. Whole-cell lysates were prepared from 1 ml of the subculture by harvesting cells through centrifugation and resuspending the cell pellets in 100 μl of SDS-loading buffer at 10x concentration. The rest of the culture was centrifuged and the supernatant (SN) containing the type III secreted proteins was filtered through a 0.45 μm-pore-size filter. Proteins were precipitated with 10% trichloroacetic acid, washed once with acetone, dried, and resuspended in 90 μl of SDS-loading buffer at 100 x concentration. Materials from 150 μl of bacterial cells and 1.5 ml of culture supernatants were analyzed by western with the indicated antibodies.

### Site-directed *in vivo* photo-crosslinking

The photoreactive amino acid *p*-benzoyl-L-phenylalanine (*p*Bpa) was incorporated into the protein of interest by chromosomally replacing the desired site-specific codon by the TAG amber codon. *S*. Typhimurium mutant strains were then co-transformed with the plasmids pSUP (encoding the orthogonal suppressor tRNA/aminoacyl-tRNA synthetase pair) and pSB3292 (encoding the HilA master regulator to strengthen SPI-1 expression). Overnight cultures were diluted 1:20 in pre-warmed 0.3 M NaCl LB supplemented with 50 μg/ml ampicillin, 10 μg/ml chloramphenicol, 1 Mm *p*Bpa, and 0.075% arabinose for 6 h. Bacterial cells were divided into two 1 ml samples, one was kept in the dark as untreated control, and the other was exposed to UV light (365 nm wavelength) for 45 min. Bacteria cells were collected by centrifugation, resuspended in 100 μl of SDS-loading buffer (10x concentration), and materials from 200 μl of bacterial cell culture were analyzed by Western-blot.

### Western-blot analysis

Protein samples were subjected to SDS-PAGE and transferred to nitrocellulose membranes. Membranes were blocked with Tris-buffered saline (TBS) containing 5% nonfat dry milk and probed with the indicated primary antibodies. Blots were then treated with DyLight 680- and DyLight 800-conjugated goat secondary antibodies directed to rabbit or mouse IgG, respectively. Detection was carried out using the Odyssey Li-Cor system and quantification of bands was performed using Image Studio Lite software (LI-COR Biosciences).

### Construction of the *orgA* random mutant library

The *orgA* gene fused to the M45-epitope tag was cloned into the pBAD24 vector^59^ and used as template for the error-prone PCR. The random mutagenesis was performed using primers *orgA*_mut_F (CCGTTTTTTTGGGCTAGCAGGAGGAATTCACCATG) and *orgA*_mut_R (GGAGGTAGGCGATCCCTACTCCGATCCATGGC) following a previously established protocol^34^, employing unbalanced dNTPs and 0.05 mM MnCl_2_ to decrease the fidelity of the Taq polymerase. The randomized PCR product was purified by gel extraction and cloned back into the pBAD24 vector using Gibson assembly. The random mutant library was then transformed into electrocompetent DH5α *E. coli* cells and a dilution was plated on ampicillin LB plates to determine the library size. Twenty individual colonies were handpicked, and Sanger sequenced to estimate the mutation rate using Mutanalyst software^60^, which was 1.6 mutations on average per sequence. The expected error spectrum biases for error-prone PCR with Taq polymerase were observed in the mutant libraries. DH5α transformed colonies were then pooled by scraping the plate with 3 ml of LB and plasmid DNA was extracted and transformed into the Δ*orgA S*. Typhimurium deletion strain and transformants were recovered on ampicillin plates. Colonies scraped from these plates constituted the input library.

### *In vivo* selection scheme for T3SS function

For each iteration, three 15 cm dishes seeded with ∼1×10^7^ Henle-407 cells were grown one day prior to invasion. The transformed *S*. Typhimurium library described above (input library) was pooled, grown under SPI-1 T3SS inducing conditions, and used as inoculum for invasion assays at a multiplicity of infection (m.o.i.) of 10 in pre-warmed Hank’s balanced salt solution (a low m.o.i. was used to diminish the extent of cooperative uptake of *Salmonella*^61^). The infection was let to proceed for 1 h and then the cells were washed 3 times with pre-warmed PBS and incubated in culture medium containing 100 μg/ml gentamicin to kill extracellular bacteria for an additional 1 h. After the gentamicin treatment, cells were washed as above and lysed with 0.1% Na-deoxycholate (in PBS). After addition of DNase I (with 100 μg/ml) to degrade the released genomic DNA from the host cells, the samples were centrifuged at 5,000 x *g* for 5 min to isolate bacteria from soluble cellular debris. The *Salmonella*-containing pellet was resuspended in LB, grown overnight, and used as the inoculum to initiate the next invasion cycle. After three cycles, intracellular *Salmonella* released from the last iteration (output library) were amplified in LB at 37°C, and plasmid DNA was extracted from the input and output libraries and processed to construct an Illumina sequencing library as follows. Four sub-amplicons that tile the length of *orgA* gene were generated, purified, and barcoded using a sample-specific index for input and output libraries. Paired-end DNA-sequencing (2 × 150 bp) was performed using an Illumina HiSeq4000 platform at the Yale Center for Genomic Analysis.

### Next generation sequencing analysis

Paired-end reads were merged with FLASH^62^, quality filtered, trimmed to the region of interest, and mapped to the *orgA* gene using Bowtie2^63^. Sequences containing gaps were filtered out. Mutations at a specific position with less than 10 observations were discarded from the analysis. Nucleotide sequences were then translated and an amino acid level matrix was generated using custom R scripts. A fitness score 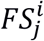 reflecting the enrichment of the amino acid substitution *i* at each position *j* relative to the wild-type residues was calculated as follows:

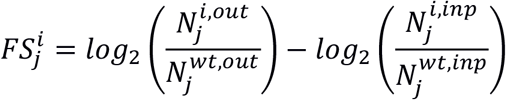

where 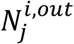 and 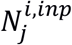 are the counts of the variant *i* in the output and input libraries, respectively, and 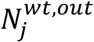 and 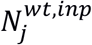 are the original wild-type amino acid counts in the output and input libraries, respectively.

### Protein structure modelling

Structural homology models of the InvC and OrgB proteins were individually generated using the I-TASSER server^64^. The first 8 residues of InvC were removed as they poorly align with the FliI monomer. OrgB was modelled as a full-length protein. The individual models were superimposed onto the solved structures of the FliI-FliH2 complex (PDB ID code 5B0O) to generate the InvC-OrgB complex or onto the EscN homohexamer structure (PDB ID code 6NJP) to obtain the InvC hexameric model. The PrgH-OrgA and OrgA-SpaO complexes were built using the AlphaFold2-Multimer framework^27^. The ‘AlphaFold2_advanced’ notebook from ColabFold was executed using default parameters^65^. MMseq2 was used for MSA generation and the models were ranked according to their pTMscore. The PDB files for the AF2 modelled 2PrgH:OrgA and OrgA:SpaO complexes are provided in S1 and S2 files, respectively. The closest residues between two different protein chains were identified using the “select” command in UCSF Chimera^66^. Structures were visualized in UCSF Chimera or UCSF ChimeraX^67^.

## Supporting information

OrgA-SpaO

2PrgH-OrgA

## DATA AVAILABILITY

All data associated with this paper are included in the submission.

## ACKNOWLEDGMENTS

JES was supported in part by the Pew Latin American Fellows Program in the Biomedical Sciences. This work was supported by Grants AI126158 (to M. L.-T.) and AI030492 (to J. E. G.) from the National Institute of Allergy and Infectious Diseases.

