## Supplementary Materials for "Assembly and architecture of the type III secretion sorting platform"

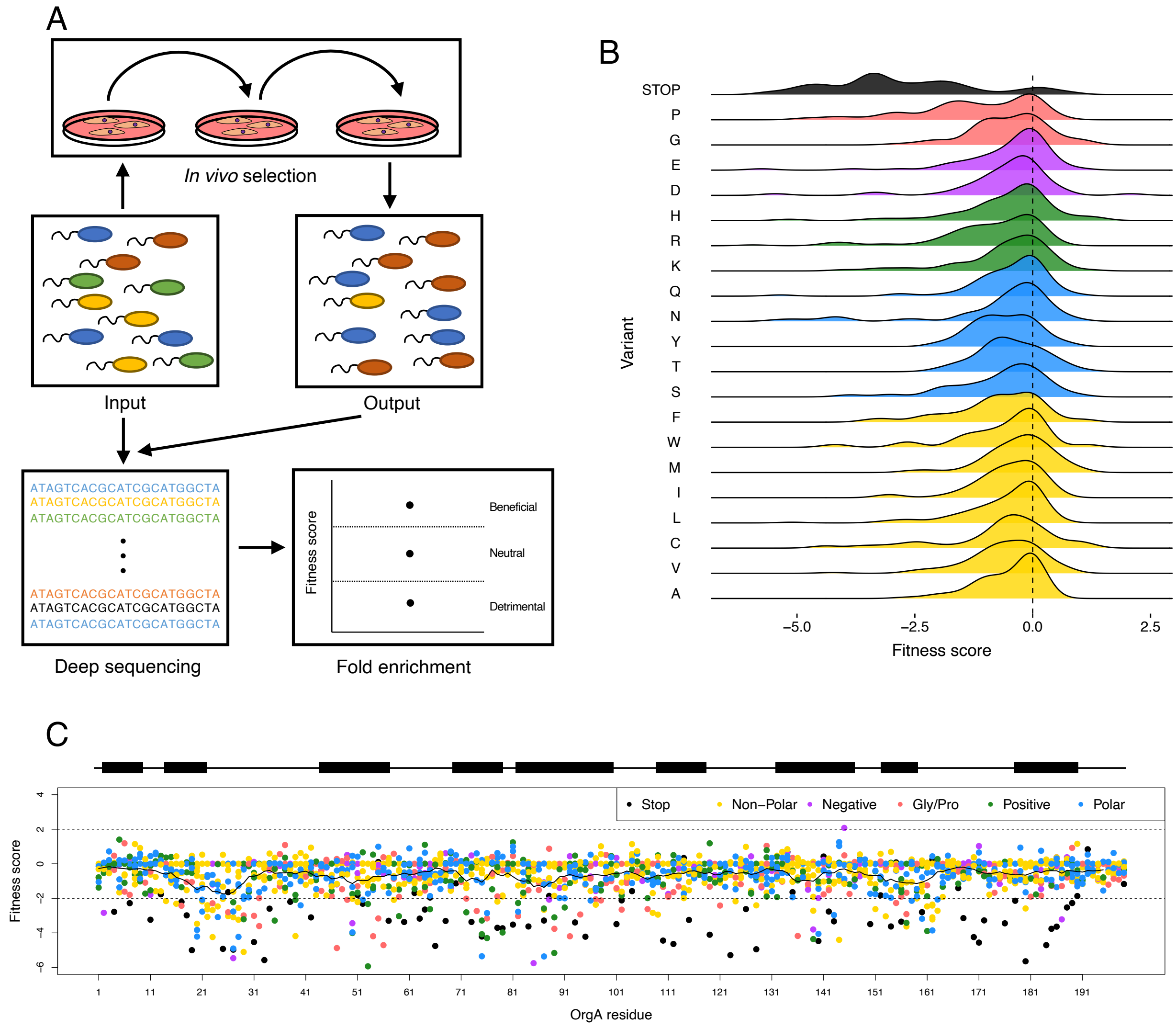


**Figure S1. Deep mutational scanning of *orgA*.** (**A**) General workflow for the deep mutational scanning. The Δ*orgA S*. Typhimurium strain transformed with a random mutant library of the *orgA* gene was used as the input library and serially subjected to 3 passages of cell culture invasion and gentamicin protection assay, a phenotype strictly dependent on OrgA function. Deep sequencing of the library before and after selection enabled the estimation of a fitness score for each variant. (**B**) Ridgeplot showing the density distribution of the fitness scores for each of the 21 possible substitutions (20 aa+stop codon). (**C**) Plot showing the fitness score of each variant per-position. Each data-point corresponds to a point mutant and is colored according to the biochemical properties of the substitution (see the upper right legend). Stop codons are colored in black. The superimposed black trace represents the moving average of the relative fitness score (rolling window size of 5 residues). Horizontal dashed lines indicate the threshold for $\log_2$ fold change > |2|. The secondary structure of the OrgA protein is shown above the plot (α-helices depicted as black rectangles).

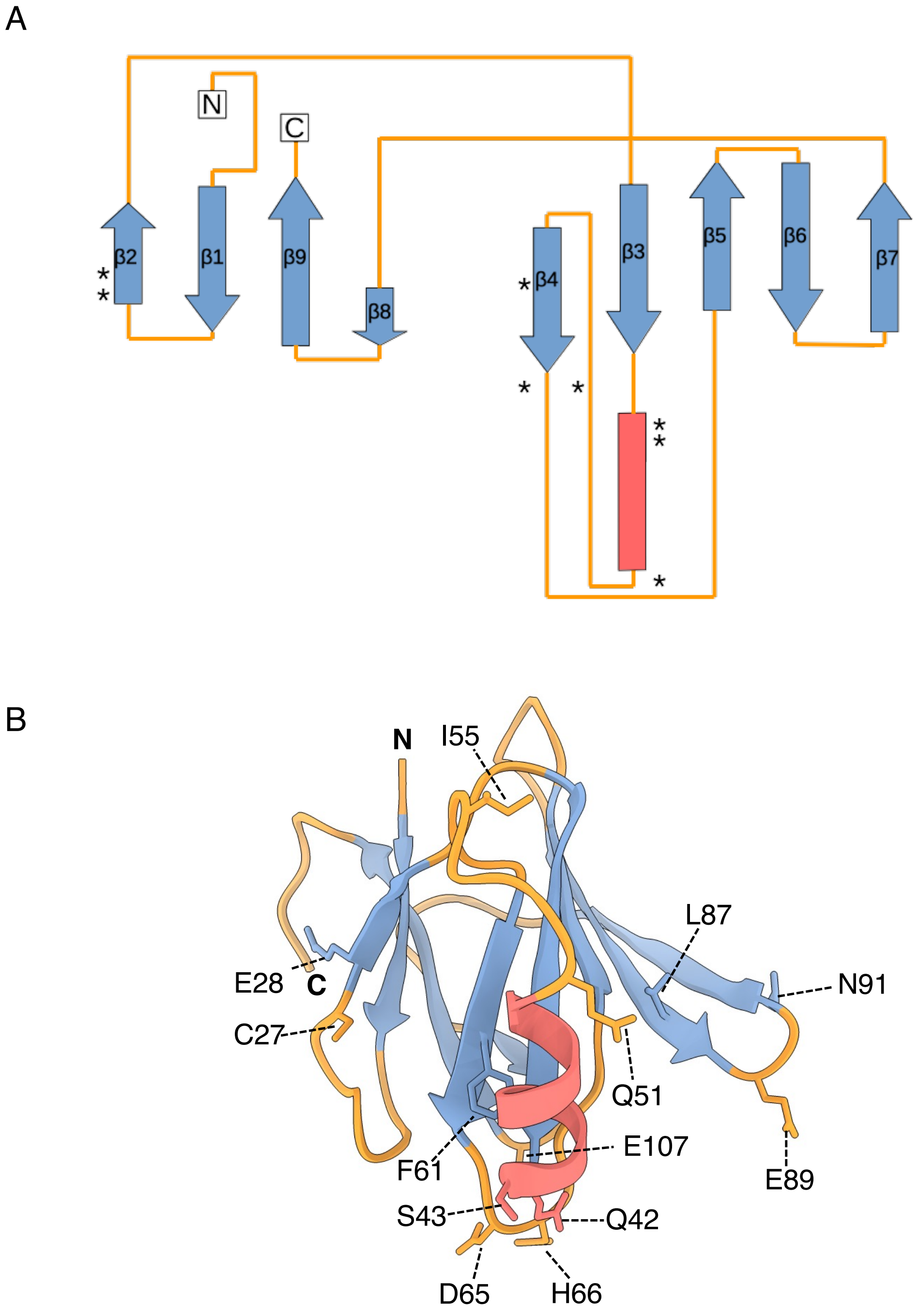


**Figure S2. Topology diagram and structure of the cytoplasmic domain of PrgH.** (**A**) 2D topology of the PrgH cytoplasmic domain. The locations of PrgH sites where introduction of *p*Bpa results in crosslinks to OrgA are indicated with an asterisk. PrgH helix, β strands, and loops are colored in red, blue, and orange, respectively. (**B**) Ribbon representation of the cytoplasmic domain of PrgH (PDB ID code: 3J1W). The residues replaced by *p*Bpa are indicated. The structure was colored to match secondary elements in A.

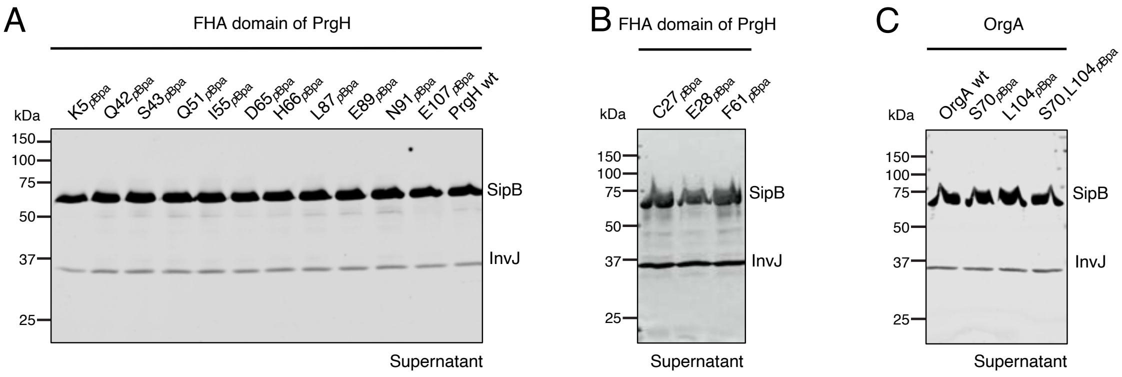


**Figure S3. Type III protein secretion profile of PrgH and OrgA *p*Bpa mutants (related to Figure 1).** Culture supernatants of *S*. Typhimurium strains expressing the indicated chromosomally-encoded *p*Bpa-containing PrgH (**A** and **B**) or OrgA (**C**) variants were analyzed by Western-blot for the presence of type III secreted proteins SipB and InvJ with specific antibodies.

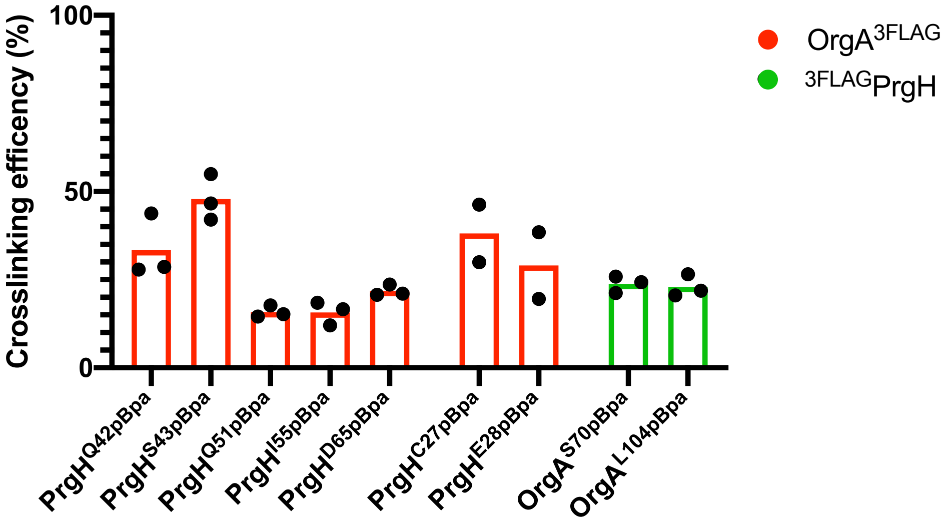


**Figure S4. Quantification of the proportion of PrgH or OrgA crosslinked species (related to Figure 1).** The proportion of the immunoblot signal corresponding to $OrgA^{3FLAG}$ (red) or $^{3FLAG}PrgH$ (green) crosslinked adduct bands on the indicated UV irradiated strain was quantified with a LICOR-Odyssey system. Bars represent the mean of three biological replicates (n=2 for $PrgH^{C27\mathit{p}Bpa}$ and $PrgH^{E28\mathit{p}Bpa}$ strains).

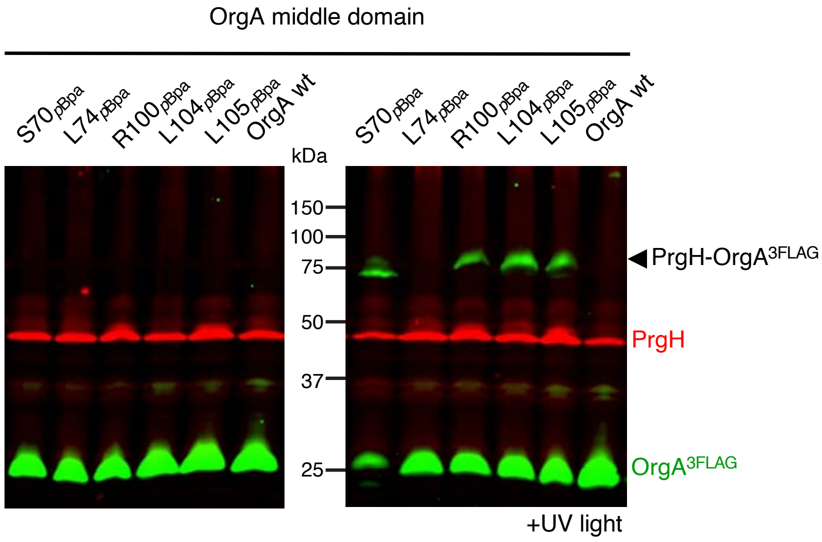


**Figure S5. Mapping of the PrgH-binding interface in OrgA.** Based on the AlphaFold2 model of the PrgH-OrgA complex (Fig 1A), *p*Bpa was introduced at the indicated positions in the middle domain of OrgA. Whole cell lysates of *S*. Typhimurium strains expressing $OrgA^{3FLAG}$ wild-type (wt) or the indicated $OrgA^{3FLAG}$ *p*Bpa variants, that were exposed to UV light or left untreated, were analyzed by immunoblot with antibodies directed to PrgH (red channel, anti-rabbit) or the FLAG epitope (green channel, anti-mouse). The crosslinked adduct that displays a mass consistent with the PrgH-$OrgA^{3FLAG}$ complex (44.4 kDa + 25.7 kDa) is indicated.

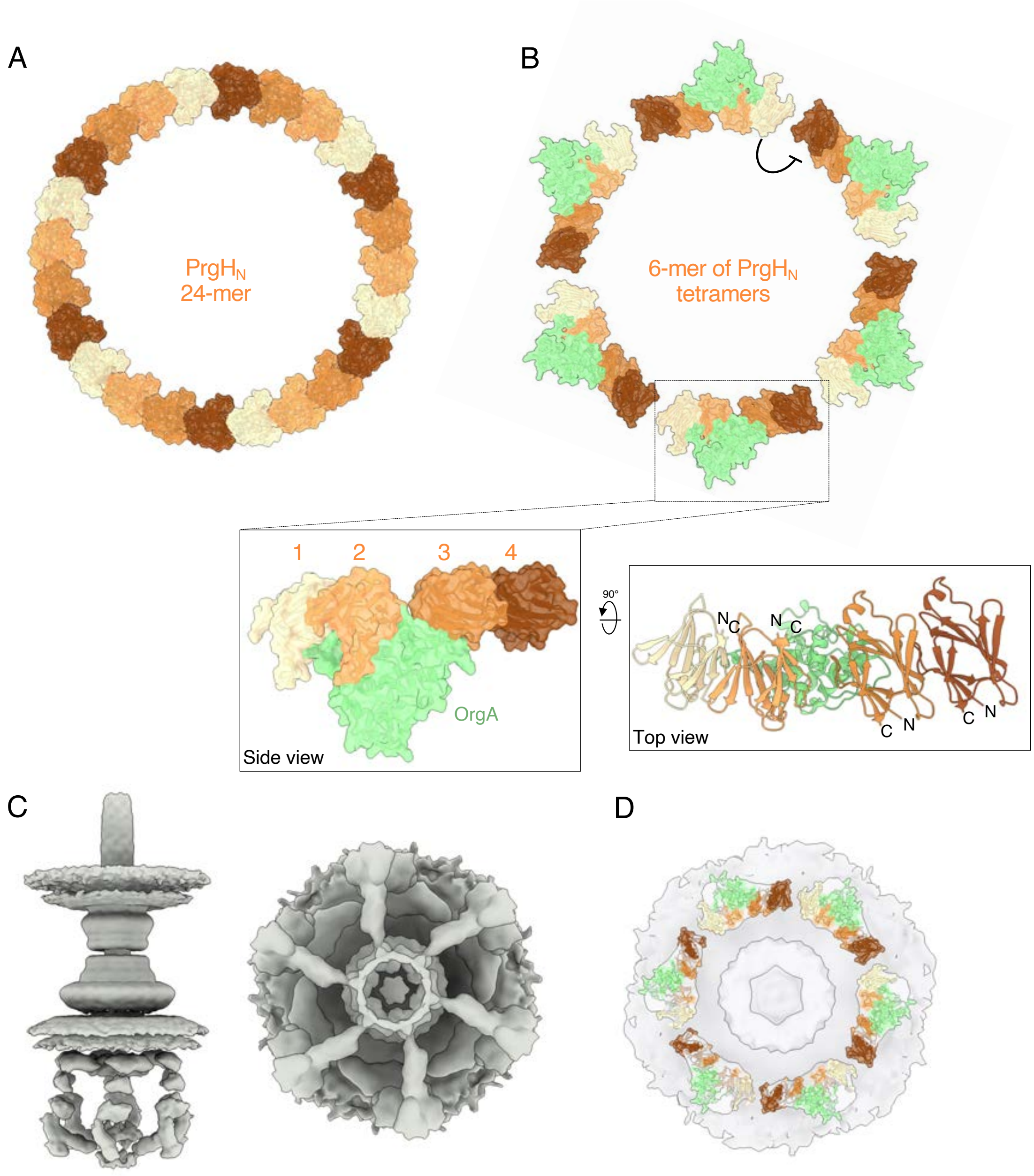


**Figure S6. Proposed mechanistic model for the OrgA-driven rearrangement of the cytosolic domain of $PrgH_N$.** (**A**) $PrgH_N$ forms a 24-unit ring when the sorting platform is absent (**B**). OrgA binding to two adjacent subunits of $PrgH_N$ (labeled as 2 and 3), results in their change of orientation relative to one another, which in turn triggers the reorganization of the immediatedly adjacent subunits (labeled 1 and 4) forming a tetrameric structure (**B**). The inset shows the side and top view of the $PrgH_N$-OrgA patch model. The N- and C-terminal ends of $PrgH_N$ are also denoted (**B**). Such spatial reorganization results in the formation of 6 discrete patches where the six-fold symmetry sorting platform is docked, shown in the front and bottom views of the cryo ET *in situ* structures of the S. Typhimurium injectisome (EMDB accession code EMD-8544) (**C**). The manual fitting of the modeled $PrgH_N$-OrgA structure onto the 6 patches of the *in situ* cryo ET structure of the injectisome is shown in (**D**).

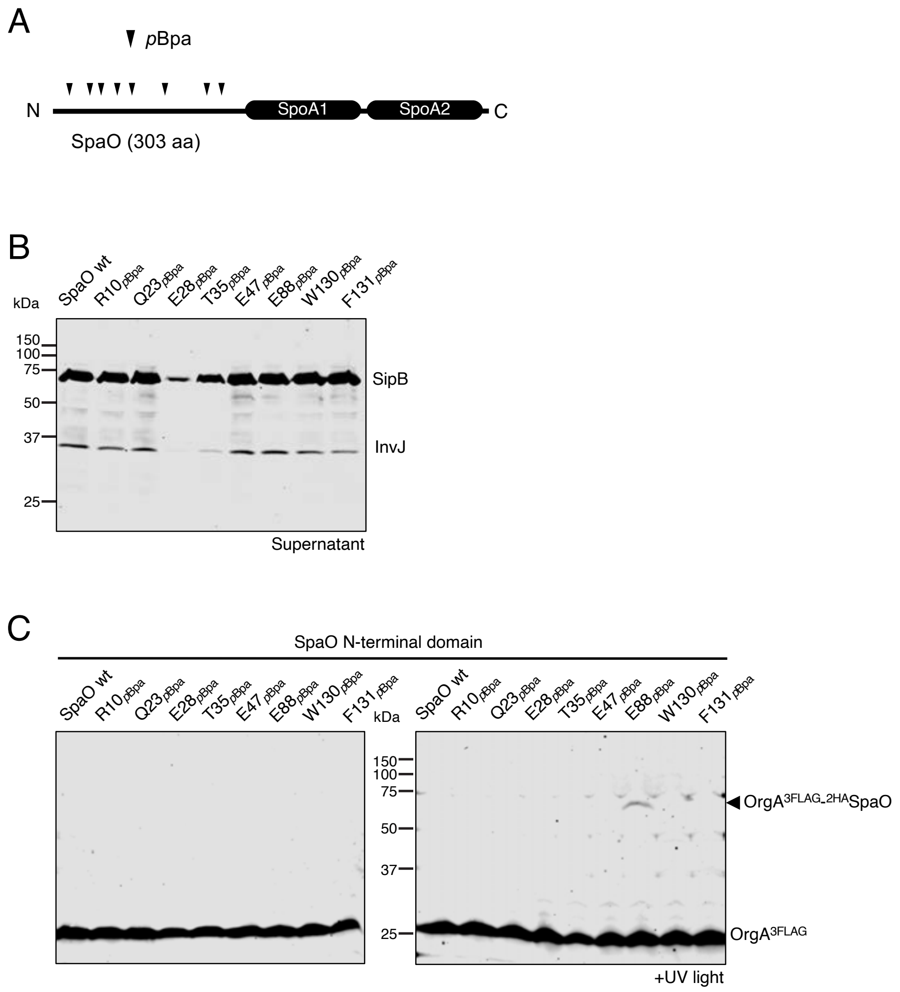


**Figure S7. Photocrosslinking survey defines the OrgA-binding domain in SpaO.** (**A**) Schematic diagram showing the domain organization of the SpaO protein. The positions where *p*Bpa was introduced is indicated by arrowheads. The location of the two SPOA domains in the SpaO C-terminal half is also indicated. (**B**) Culture supernatants of *S*. Typhimurium chromosomally expressing OrgA$^{3FLAG}$ and $^{2HA}$SpaO wild-type (wt) or the indicated $^{2HA}$SpaO *p*Bpa-containing variant were analyzed by Western blot for the presence of the type III secreted proteins SipB and InvJ. (**C**) Survey of the N-terminal half of SpaO for amino acids that can crosslink with OrgA. Whole cell lysates of *S*. Typhimurium strains expressing the OrgA$^{3FLAG}$ and the wild-type $^{2HA}$SpaO (wt) or the indicated $^{2HA}$SpaO *p*Bpa variants after exposure to UV light or left untreated. Lysates were analyzed by immunoblot using antibodies directed to the FLAG epitope. The detected crosslink between $^{2HA}$SpaO$^{E88pBpa}$ and OrgA$^{3FLAG}$ is indicated.

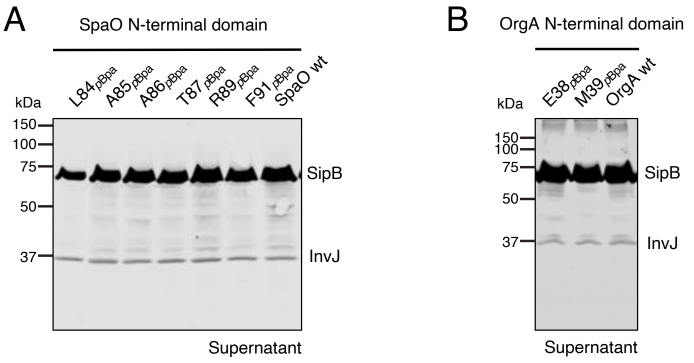


**Figure S8. Type III protein secretion profile of SpaO and OrgA *p*Bpa mutants (related to Figure 2).** Culture supernatants of *S*. Typhimurium strains chromosomally expressing $\text{OrgA}^{3FLAG}$ (**A**) or $^{2HA}$SpaO (**B**) and the indicated *p*Bpa-containing $^{2HA}$SpaO (A) or $\text{OrgA}^{3FLAG}$ (B) variant, were analyzed by Western blot for the presence of the type III secreted proteins SipB and InvJ.

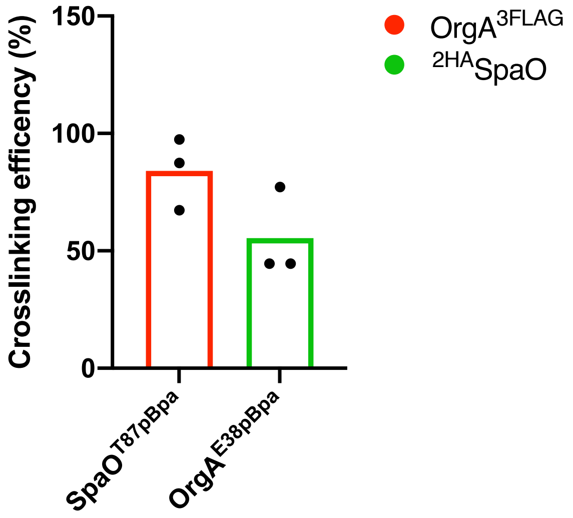


**Figure S9. Quantification of the proportion of crosslinked species (related to Figure 2).** The proportion of the immunoblot signal corresponding to OrgA$^{3FLAG}$ (red) or $^{2HA}$SpaO (green) crosslinked adduct bands on the indicated UV irradiated strains was quantified with a LICOR-Odyssey system. Bars represent the mean of the three biological replicates.

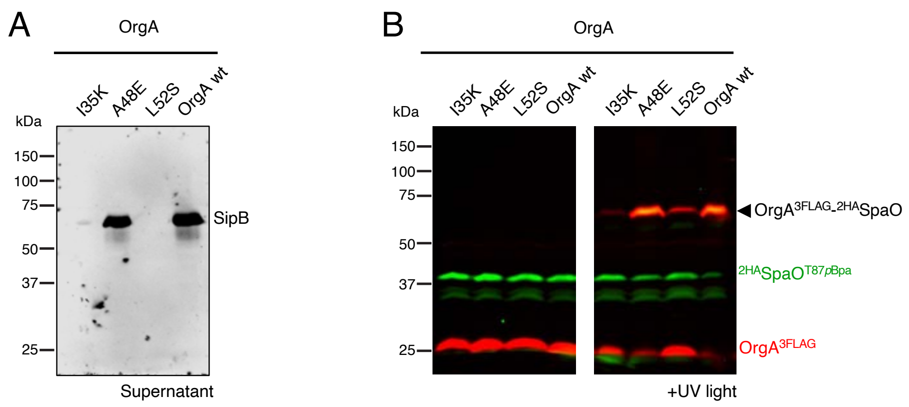


**Figure S10. Effect of OrgA mutations on its interaction with SpaO.** (**A**) Culture supernatants of the *S*. Typhimurium strains chromosomally expressing $^{2HA}$SpaO$^{T87pBpa}$ and wild-type OrgA$^{3FLAG}$ (wt) or the indicated OrgA$^{3FLAG}$ point mutant were analyzed by Western blot for the presence of the type III secreted protein SipB. (**B**) Whole-cell lysates of the indicated strains after exposure to UV light or left untreated. Lysates were analyzed by Western blot with antibodies directed to the FLAG (red channel, anti-rabbit) or HA (green channel, anti-mouse) epitopes. The detected crosslinks between $^{2HA}$SpaO and OrgA$^{3FLAG}$ are indicated.

A

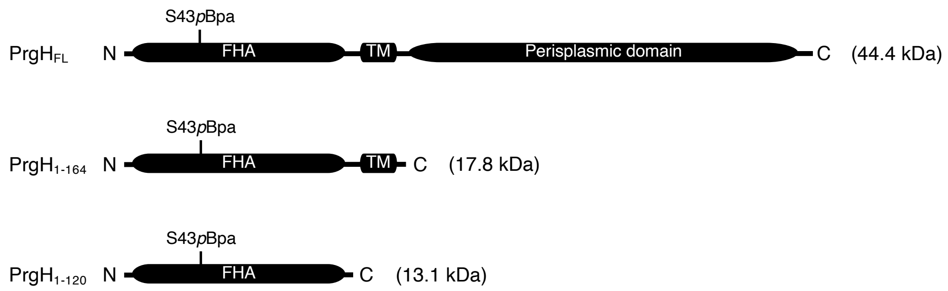


B

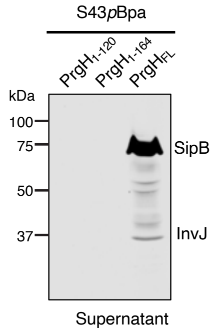


**Figure S11. OrgA has affinity for the FHA domain of PrgH but formation of higher-order structures requires PrgH ring assembly (related to Figure 3D).** (**A**) Domain organization of PrgH and schematics of the different truncated versions used in this study. The localization of the *p*Bpa residue is indicated. FL, full-length; TMH, transmembrane helix; FHA, forkhead-associated domain. (**B**) Culture supernatants of the *S*. Typhimurium strains encoding the indicated $PrgH^{S43pBpa}$ allele were analyzed by Western blot for the presence of the type III secreted proteins SipB and InvJ.

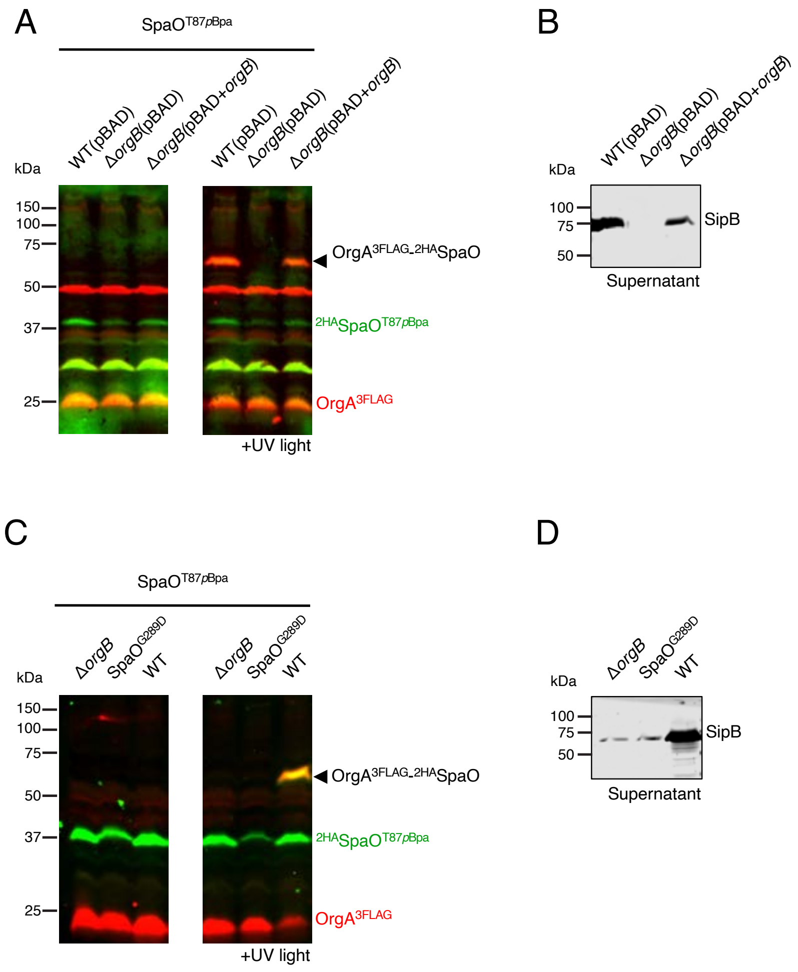


**Fig S12. The OrgB protein enables the formation of the SpaO-OrgA complex.** (**A**) Whole-cell lysates of *S*. Typhimurium strains chromosomally encoding the $^{2HA}$SpaO$^{T87\textit{p}Bpa}$ allele in the context of the wild-type or the Δ*orgB* mutant strain, carrying an empty pBAD24 vector (pBAD) or the same plasmid backbone expressing *orgB* (pBAD+*orgB*) (as indicated) after exposure to UV light or left untreated. Cell lysates were analyzed by Western blot with antibodies directed to the FLAG (red channel, anti-rabbit) or HA (green channel, anti-mouse) epitopes. (**B**) The culture supernatants of the indicated *S*. Typhimurium strains were analyzed by Western blot for the presence of the type III secreted protein SipB. (**C**) *S*. Typhimurium strains encoding the $^{2HA}$SpaO$^{T87\textit{p}Bpa}$ allele in the Δ*orgB*, SpaO$^{G289D}$, or wild-type (WT) background were exposed to UV light or left untreated as indicated and analyzed by Western blot with antibodies to the FLAG (red) or HA (green) epitopes. (**D**) The culture supernatants of the indicated *S*. Typhimurium strains were analyzed by Western blot for the presence of the type III secreted protein SipB.

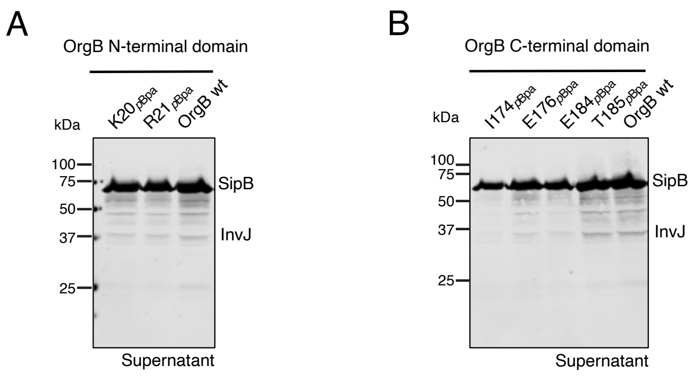


**Figure S13. Type III protein secretion profile of OrgB *p*Bpa mutants (related to Figure 4).** Culture supernatants of *S*. Typhimurium strains expressing $^{3FLAG}$SpaO (**A**) or InvC$^{3FLAG}$ (**B**), and the indicated OrgB$^{M45}$ mutants carrying *p*Bpa at its N- (**A**) or C-terminal (**B**) regions were analyzed by Western blot for the presence of the type III secreted proteins SipB and InvJ.

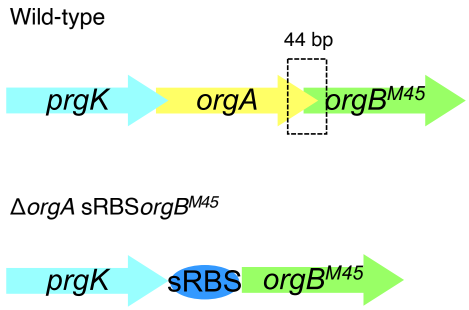


**Fig S14. Schematic representation of the strategy used to uncouple OrgB synthesis from *orgA* translation (related to figure 5C).** Schematic representation of the upstream genetic organization of the *orgB* gene. The *orgA* gene was replaced by a synthetic RBS (sRBS) to uncouple OrgB synthesis from *orgA* translation.

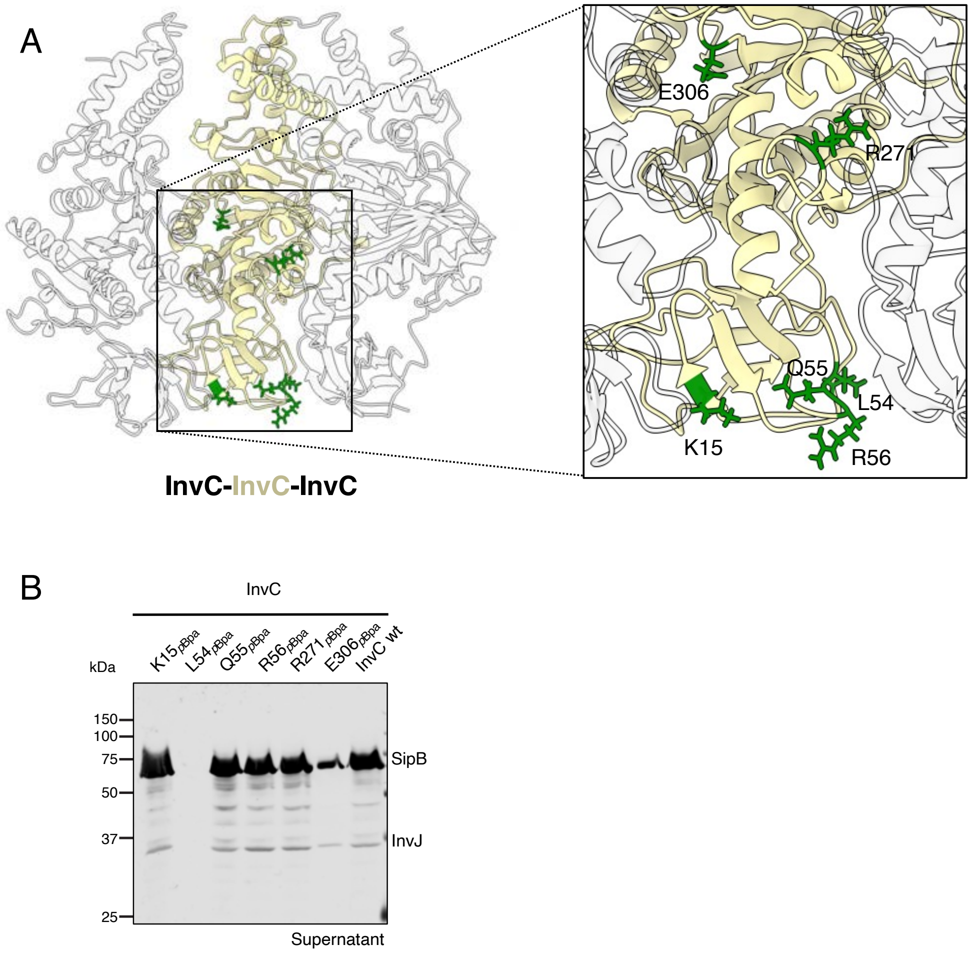


**Figure S15. Identification of residues at the interface between InvC molecules within its hexameric context (related to figure 5E).** (**A**) Ribbon representation of the model for the InvC hexameric structure. For clarity, only a single InvC subunit (colored in light yellow) flanked by two neighboring subunits (in white) is shown. The InvC residues chosen for *p*Bpa incorporation are shown in in green. Inset shows a close-up view of the location of the InvC residues replaced by *p*Bpa. (**B**) Culture supernatants of *S*. Typhimurium strains expressing the indicated *p*Bpa mutant of InvC$^{3FLAG}$ were analyzed by Western blot for the presence of the type III secreted proteins SipB and InvJ.

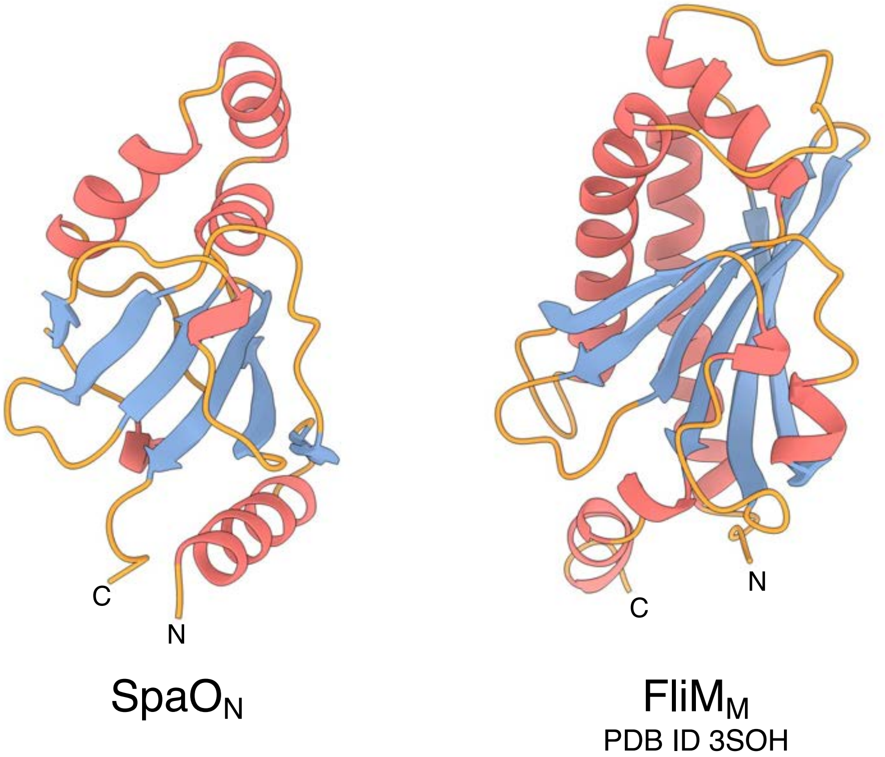


**Figure S16. $SpaO_N$ shares structural similarity with $FliM_M$.** Ribbon representation of the AlphaFold 2 model of the N-terminal domain of SpaO (residues 10-138) and the solved structure of the middle domain of FliM (residues 46-233, PDB ID code: 3SOH, chain A) from *Thermotoga maritima*. Helices, $\beta$ strands, and loops are colored in red, blue, and orange, respectively.

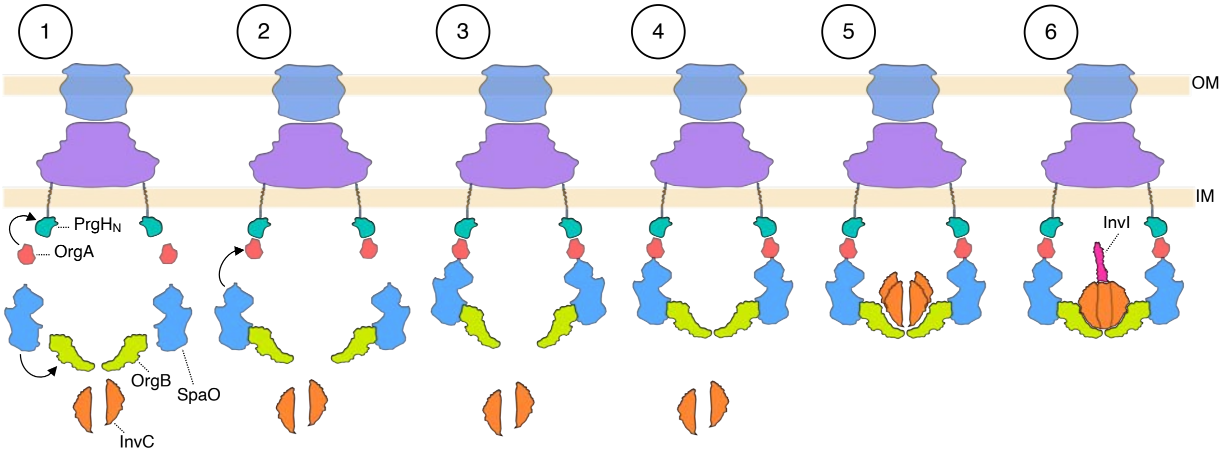


**Fig S17. Proposed pathway for the assembly of the *Salmonella* sorting platform.** (1 through 3) OrgA docks onto PrgH$_N$ concomitantly to PrgH ring assembly. OrgB engages the C-terminal region of SpaO triggering conformational change that make it permissive for binding to PrgH-bound OrgA and completing the assemble of the pods. (4 and 5) Assemble of the OrgB craddle provides a structural scaffold to recruit InvC, facilitating its oligomerization. (6) Incorporation of InvI onto the InvC hexamer results in a functional type III secretion sorting platform.

**Table S1. Unified nomenclature for relevant structural components of T3SSs in representative bacteria and flagellar homologues.**

| **Function** | **Unified Sct** | ***Salmonella* SPI-1** | ***Salmonella* SPI-2** | ***Shigella*** | ***Yersinia*** | **Flagella** |
|---|---|---|---|---|---|---|
| OM secretin | SctC | InvG | SsaC | MxiD | YscC | - |
| IM ring | SctD | PrgH | SsaD | MxiG | YscD | - |
| IM ring | SctJ | PrgK | SsaJ | MxiJ | YscJ | FliF |
| Linker protein | SctK | OrgA | STM1410 | MxiK | YscK | FliG |
| Major sorting platform | SctQ | SpaO | SsaQ | Spa33 | YscQ | FliM/FliN |
| Spokes | SctL | OrgB | SsaK | MxiN | YscL | FliH |
| ATPase | SctN | InvC | SsaN | Spa47 | YscN | FliI |
| Central stalk | SctO | InvI | SsaO | Spa13 | YscO | FliJ |

**Table S2. Strains and plasmids used in this study.**

| Strain | Relevant genotype | Source or reference |
|---|---|---|
| SB300 | Wild-type *Salmonella* | PMID: 7015147 |
| SB1677 | Δ*orgA* | J. Kato (unpublished) |
| SB1681 | $OrgA^{3FLAG}$ | PMID: 21292939 |
| SB3855 | $OrgA^{3FLAG}$, PrgH Q42 to TAG | This study |
| SB3851 | $OrgA^{3FLAG}$, PrgH S43 to TAG | This study |
| SB3760 | $OrgA^{3FLAG}$, PrgH Q51 to TAG | This study |
| SB3856 | $OrgA^{3FLAG}$, PrgH I55 to TAG | This study |
| SB3857 | $OrgA^{3FLAG}$, PrgH D65 to TAG | This study |
| SB3858 | $OrgA^{3FLAG}$, PrgH H66 to TAG | This study |
| SB3852 | $OrgA^{3FLAG}$, PrgH L87 to TAG | This study |
| SB3853 | $OrgA^{3FLAG}$, PrgH E89 to TAG | This study |
| SB3854 | $OrgA^{3FLAG}$, PrgH N91 to TAG | This study |
| SB3859 | $OrgA^{3FLAG}$, PrgH E107 to TAG | This study |
| SB3938 | $OrgA^{3FLAG}$, PrgH C27 to TAG | This study |
| SB4146 | $OrgA^{3FLAG}$, PrgH E28 to TAG | This study |
| SB4147 | $OrgA^{3FLAG}$, PrgH F61 to TAG | This study |
| SB3947 | $OrgA^{3FLAG}$, PrgH S43 to TAG P121 to TAA | This study |
| SB3948 | $OrgA^{3FLAG}$, PrgH S43 to TAG S165 to TAA | This study |
| SB3888 | $OrgA^{3FLAG}$, $^{2HA}SpaO$ | This study |
| SB3940 | $OrgA^{3FLAG}$, $^{2HA}SpaO$ L84 to TAG | This study |

| | | |
|---|---|---|
| SB3941 | OrgA$^{3FLAG}$, $^{2HA}$SpaO A85 to TAG | This study |
| SB3942 | OrgA$^{3FLAG}$, $^{2HA}$SpaO A86 to TAG | This study |
| SB3943 | OrgA$^{3FLAG}$, $^{2HA}$SpaO T87 to TAG | This study |
| SB3944 | OrgA$^{3FLAG}$, $^{2HA}$SpaO R89 to TAG | This study |
| SB3945 | OrgA$^{3FLAG}$, $^{2HA}$SpaO F91 to TAG | This study |
| SB3934 | OrgA$^{3FLAG}$, $^{2HA}$SpaO R10 to TAG | This study |
| SB3935 | OrgA$^{3FLAG}$, $^{2HA}$SpaO Q23 to TAG | This study |
| SB3936 | OrgA$^{3FLAG}$, $^{2HA}$SpaO E28 to TAG | This study |
| SB3937 | OrgA$^{3FLAG}$, $^{2HA}$SpaO T35 to TAG | This study |
| SB3890 | OrgA$^{3FLAG}$, $^{2HA}$SpaO E47 to TAG | This study |
| SB3931 | OrgA$^{3FLAG}$, $^{2HA}$SpaO E88 to TAG | This study |
| SB3932 | OrgA$^{3FLAG}$, $^{2HA}$SpaO W130 to TAG | This study |

| | | |
|---|---|---|
| SB3933 | OrgA$^{3FLAG}$, $^{2HA}$SpaO F131 to TAG | This study |
| SB4167 | OrgA$^{M45}$, $^{3FLAG}$PrgH | This study |
| SB4158 | OrgA$^{M45}$ S70 to TAG, $^{3FLAG}$PrgH | This study |
| SB4159 | OrgA$^{M45}$ L104 to TAG, $^{3FLAG}$PrgH | This study |
| SB4160 | OrgA$^{M45}$ S70 to TAG and L104 to TAG, $^{3FLAG}$PrgH | This study |
| SB4056 | OrgA$^{3FLAG}$ I35K, $^{2HA}$SpaO T87 to TAG | This study |
| SB4058 | OrgA$^{3FLAG}$ A48E, $^{2HA}$SpaO T87 to TAG | This study |
| SB4059 | OrgA$^{3FLAG}$ L52S, $^{2HA}$SpaO T87 to TAG | This study |
| SB3960 | OrgA$^{3FLAG}$, $^{2HA}$SpaO T87 to TAG and G289D | This study |
| SB4037 | OrgA$^{3FLAG}$ E38 to TAG, $^{2HA}$SpaO | This study |
| SB4038 | OrgA$^{3FLAG}$ M39 to TAG, $^{2HA}$SpaO | This study |
| SB2691 | OrgB$^{M45}$, $^{3FLAG}$SpaO | PMID: 30668610 |
| SB3965 | OrgB$^{M45}$ K20 to TAG, $^{3FLAG}$SpaO | This study |

| | | |
|---|---|---|
| SB3966 | $OrgB^{M45}$ R21 to TAG, $^{3FLAG}SpaO$ | This study |
| SB3971 | $OrgB^{M45}$, $InvC^{3FLAG}$ | This study |
| SB3967 | $OrgB^{M45}$ I174 to TAG, $InvC^{3FLAG}$ | This study |
| SB3968 | $OrgB^{M45}$ E176 to TAG, $InvC^{3FLAG}$ | This study |
| SB3969 | $OrgB^{M45}$ E184 to TAG, $InvC^{3FLAG}$ | This study |
| SB3970 | $OrgB^{M45}$ T185 to TAG, $InvC^{3FLAG}$ | This study |
| SB1680 | $InvC^{3FLAG}$ | PMID: 21292939 |
| SB4019 | $InvC^{3FLAG}$ K15 to TAG | This study |
| SB4020 | $InvC^{3FLAG}$ L54 to TAG | This study |
| SB4021 | $InvC^{3FLAG}$ Q55 to TAG | This study |
| SB4022 | $InvC^{3FLAG}$ R56 to TAG | This study |
| SB4016 | $InvC^{3FLAG}$ R271 to TAG | This study |
| SB4017 | $InvC^{3FLAG}$ E306 to TAG | This study |
| SB3961 | PrgH S43 to TAG, $OrgA^{3FLAG}$, $^{2HA}SpaO$ T87 to TAG | This study |
| SB4144 | $OrgA^{3FLAG}$, $OrgB^{M45}$, $^{2HA}SpaO$ N285 to TAG | This study |
| SB4142 | $OrgA^{3FLAG}$, $OrgB^{M45}$, $^{2HA}SpaO$ T87 to TAG and N285 to TAG | This study |
| SB3861 | Δ*orgA*, PrgH S43 to TAG | This study |

| | | |
|---|---|---|
| SB3873 | Δ*orgB*, $OrgA^{3FLAG}$, PrgH S43 to TAG | This study |
| SB3868 | Δ*spaO*, $OrgA^{3FLAG}$, PrgH S43 to TAG | This study |
| SB3877 | Δ*invC*, $OrgA^{3FLAG}$, PrgH S43 to TAG | This study |
| SB3878 | Δ*invI*, $OrgA^{3FLAG}$, PrgH S43 to TAG | This study |
| SB3880 | PrgK M1 to ATA, $OrgA^{3FLAG}$, PrgH S43 to TAG | This study |
| SB3875 | Δ*invG*, $OrgA^{3FLAG}$, PrgH S43 to TAG | This study |
| SB3879 | Δ*spaPQRS*, $OrgA^{3FLAG}$, PrgH S43 to TAG | This study |
| SB3952 | Δ*orgA*, $^{2HA}$SpaO T87 to TAG | This study |
| SB3950 | Δ$prgH_N$, $OrgA^{3FLAG}$, $^{2HA}$SpaO T87 to TAG | This study |
| SB3949 | Δ*orgB*, $OrgA^{3FLAG}$, $^{2HA}$SpaO T87 to TAG | This study |
| SB3953 | Δ*invC*, $OrgA^{3FLAG}$, $^{2HA}$SpaO T87 to TAG | This study |
| SB3954 | Δ*invI*, $OrgA^{3FLAG}$, $^{2HA}$SpaO T87 to TAG | This study |
| SB3972 | Δ$prgH_N$, $OrgB^{M45}$ K20 to TAG, $^{3FLAG}$SpaO | This study |

| | | |
|---|---|---|
| SB3976 | Δ*orgA*, $OrgB^{M45}$ K20 to TAG, $^{3FLAG}SpaO$ | This study |
| SB3973 | Δ*spaO*, $OrgB^{M45}$ K20 to TAG | This study |
| SB3974 | Δ*invC*, $OrgB^{M45}$ K20 to TAG, $^{3FLAG}SpaO$ | This study |
| SB3975 | Δ*invI*, $OrgB^{M45}$ K20 to TAG, $^{3FLAG}SpaO$ | This study |
| SB4073 | *orgA::sRBS*, $OrgB^{M45}$ K20 to TAG, $^{3FLAG}SpaO$ | This study |
| SB3979 | Δ$prgH_N$, $OrgB^{M45}$ T185 to TAG, $InvC^{3FLAG}$ | This study |
| SB3980 | Δ*orgA*, $OrgB^{M45}$ T185 to TAG, $InvC^{3FLAG}$ | This study |
| SB4012 | Δ*spaO*, $OrgB^{M45}$ T185 to TAG, $InvC^{3FLAG}$ | This study |
| SB3981 | Δ*invC*, $OrgB^{M45}$ T185 to TAG | This study |
| SB4014 | Δ*invI*, $OrgB^{M45}$ T185 to TAG, $InvC^{3FLAG}$ | This study |
| SB4030 | Δ$prgH_N$, $InvC^{3FLAG}$ R56 to TAG | This study |
| SB4031 | Δ*orgA*, $InvC^{3FLAG}$ R56 to TAG | This study |
| SB4027 | Δ*orgB*, $InvC^{3FLAG}$ R56 to TAG | This study |
| SB4032 | Δ*spaO*, $InvC^{3FLAG}$ R56 to TAG | This study |
| SB4033 | Δ*invI*, $InvC^{3FLAG}$ R56 to TAG | This study |
| *E. coli* β-2163 Δnic35 | | PMID: 15748991 |

| Plasmid | Relevant genotype | Source or reference |
|---|---|---|
| pSB890 | R6K origin, suicide plasmid counter selectable with sucrose | PMID: 7997169 |
| pSB3492 | pSB890-based plasmid to introduce *orgA* clean deletion | J. Kato (unpublished) |
| pSB3489 | pSB890-based plasmid to introduce $orgA^{3FLAG}$ | PMID: 21292939 |
| pSB6124 | pSB890-based plasmid to introduce PrgH Q42 to TAG | This study |
| pSB6125 | pSB890-based plasmid to introduce PrgH S43 to TAG | This study |
| PSB6126 | pSB890-based plasmid to introduce PrgH Q51 to TAG | This study |
| PSB6127 | pSB890-based plasmid to introduce PrgH I55 to TAG | This study |
| pSB6128 | pSB890-based plasmid to introduce PrgH D65 to TAG | This study |
| pSB6038 | pSB890-based plasmid to introduce PrgH H66 to TAG | This study |
| pSB6039 | pSB890-based plasmid to introduce PrgH L87 to TAG | This study |
| pSB6040 | pSB890-based plasmid to introduce PrgH E89 to TAG | This study |

| | | |
|---|---|---|
| pSB6121 | pSB890-based plasmid to introduce PrgH N91 to TAG | This study |
| pSB6122 | pSB890-based plasmid to introduce PrgH E107 to TAG | This study |
| pSB6142 | pSB890-based plasmid to introduce PrgH C27 to TAG | This study |
| pSB6563 | pSB890-based plasmid to introduce PrgH E28 to TAG | This study |
| pSB6564 | pSB890-based plasmid to introduce PrgH F61 to TAG | This study |
| pSB6158 | pSB890-based plasmid to introduce PrgH S43 to TAG and P121 to TAA | This study |
| pSB6159 | pSB890-based plasmid to introduce PrgH S43 to TAG and S165 to TAA | This study |
| pSB6140 | pSB890-based plasmid to introduce $^{2HA}$SpaO | This study |
| pSB6151 | pSB890-based plasmid to introduce $^{2HA}$SpaO L84 to TAG | This study |
| pSB6152 | pSB890-based plasmid to introduce $^{2HA}$SpaO A85 to TAG | This study |

| | | |
|---|---|---|
| pSB6153 | pSB890-based plasmid to introduce $^{2HA}$SpaO A86 to TAG | This study |
| pSB6154 | pSB890-based plasmid to introduce $^{2HA}$SpaO T87 to TAG | This study |
| pSB6155 | pSB890-based plasmid to introduce $^{2HA}$SpaO R89 to TAG | This study |
| pSB6156 | pSB890-based plasmid to introduce $^{2HA}$SpaO F91 to TAG | This study |
| pSB6143 | pSB890-based plasmid to introduce $^{2HA}$SpaO R10 to TAG | This study |
| pSB6144 | pSB890-based plasmid to introduce $^{2HA}$SpaO Q23 to TAG | This study |
| pSB6145 | pSB890-based plasmid to introduce $^{2HA}$SpaO E28 to TAG | This study |
| pSB6146 | pSB890-based plasmid to introduce $^{2HA}$SpaO T35 to TAG | This study |

| | | |
|---|---|---|
| pSB6147 | pSB890-based plasmid to introduce $^{2HA}$SpaO E47 to TAG | This study |
| pSB6148 | pSB890-based plasmid to introduce $^{2HA}$SpaO E88 to TAG | This study |
| pSB6149 | pSB890-based plasmid to introduce $^{2HA}$SpaO W130 to TAG | This study |
| pSB6150 | pSB890-based plasmid to introduce $^{2HA}$SpaO F131 to TAG | This study |
| pSB6381 | pSB890-based plasmid to introduce OrgA$^{3FLAG}$ I35K | This study |
| pSB6383 | pSB890-based plasmid to introduce OrgA$^{3FLAG}$ A48E | This study |
| pSB6384 | pSB890-based plasmid to introduce OrgA$^{3FLAG}$ L52S | This study |
| pSB6326 | pSB890-based plasmid to introduce $^{2HA}$SpaO G289D and T87 to TAG | This study |
| pSB6378 | pSB890-based plasmid to introduce orgA$^{3FLAG}$ E38 to TAG | This study |

| | | |
|---|---|---|
| pSB6379 | pSB890-based plasmid to introduce orgA$^{3FLAG}$ M39 to TAG | This study |
| pSB4870 | pSB890-based plasmid to introduce orgB$^{M45}$ | PMID: 30668610 |
| pSB6327 | pSB890-based plasmid to introduce orgB$^{M45}$ K20 to TAG | This study |
| pSB6328 | pSB890-based plasmid to introduce orgB$^{M45}$ R21 to TAG | This study |
| pSB6329 | pSB890-based plasmid to introduce orgB$^{M45}$ I174 to TAG | This study |
| pSB6330 | pSB890-based plasmid to introduce orgB$^{M45}$ E176 to TAG | This study |
| pSB6331 | pSB890-based plasmid to introduce orgB$^{M45}$ E184 to TAG | This study |
| pSB6332 | pSB890-based plasmid to introduce orgB$^{M45}$ T185 to TAG | This study |
| pSB6371 | pSB890-based plasmid to introduce invC$^{3FLAG}$ K15 to TAG | This study |

| | | |
|---|---|---|
| pSB6372 | pSB890-based plasmid to introduce invC$^{3FLAG}$ L54 to TAG | This study |
| pSB6373 | pSB890-based plasmid to introduce invC$^{3FLAG}$ Q55 to TAG | This study |
| pSB6374 | pSB890-based plasmid to introduce invC$^{3FLAG}$ R56 to TAG | This study |
| pSB6338 | pSB890-based plasmid to introduce invC$^{3FLAG}$ R271 to TAG | This study |
| pSB6339 | pSB890-based plasmid to introduce invC$^{3FLAG}$ E306 to TAG | This study |
| pSB6395 | pSB890-based plasmid to introduce $^{2HA}$SpaO T87 to TAG and N285 to TAG | This study |
| pSB6396 | pSB890-based plasmid to introduce $^{2HA}$SpaO N285 to TAG | This study |
| pSB5144 | pSB890-based plasmid to introduce a clean deletion of *prgH*$_N$ | This study |

| | | |
|---|---|---|
| pSB6130 | pSB890-based plasmid to introduce a clean deletion of *orgB* in orgA$^{3FLAG}$ | This study |
| pSB6334 | pSB890-based plasmid to introduce a clean deletion of *orgA* in orgB$^{M45\ K20pBpa}$ | This study |
| pSB6335 | pSB890-based plasmid to introduce a clean deletion of *orgA* in orgB$^{M45\ T185pBpa}$ | This study |
| pSB6393 | pSB890-based plasmid to replace *orgA* by a synthetic RBS | This study |
| pSB3493 | pSB890-based plasmid to introduce a clean deletion of *orgB* | J. Kato (unpublished) |
| pSB3741 | pSB890-based plasmid to introduce a clean deletion of *spaO* | PMID: 21292939 |
| pSB2396 | pSB890-based plasmid to introduce a clean deletion of *invC* | PMID: 15060043 |
| pSB3749 | pSB890-based plasmid to introduce a clean deletion of *invI* | M. Lara-Tejero (unpublished) |

| | | |
|---|---|---|
| pSB376 | pSB890-based plasmid to introduce a clean deletion of *invG* | PMID: 7997169 |
| pSB3320 | pSB890-based plasmid to introduce a clean deletion of *spaPQRS* | M. Lara-Tejero (unpublished) |
| pSB6132 | pSB890-based plasmid to introduce *prgK* M1 to ATA | This study |
| pSB6336 | pSB890-based plasmid to introduce a clean deletion of *invI* in $invC^{3FLAG}$ | This study |
| pSUP-Bpa | Suppressor plasmid to incorporate pBpa into the amber codon | PMID: 16554830 |
| pSB3292 | pBAD24-hilA | PMID: 21292939 |
| pSB3785 | $pBAD\text{-}orgA^{M45}$ | M. Lara-Tejero (unpublished) |
| pSB6325 | $pBAD\text{-}orgB^{M45}$ | This study |
